# Sequestration to lipid droplets promotes histone availability by preventing turnover of excess histones

**DOI:** 10.1101/2020.08.06.237164

**Authors:** Roxan A. Stephenson, Jonathon M. Thomalla, Lili Chen, Petra Kolkhof, Mathias Beller, Michael A. Welte

## Abstract

Because dearth and overabundance of histones result in cellular defects, histone synthesis and demand are typically tightly coupled. In *Drosophila* embryos, histones H2B/H2A/H2Av accumulate on lipid droplets (LDs), cytoplasmic fat storage organelles. Without this binding, maternally provided H2B/H2A/H2Av are absent; however, the molecular basis of how LDs ensure histone storage is unclear. Using quantitative imaging, we uncover when during oogenesis these histones accumulate, and which step of accumulation is LD-dependent. LDs originate in nurse cells and are transported to the oocyte. Although H2Av accumulates on LDs in nurse cells, the majority of the final H2Av pool is synthesized in oocytes. LDs promote intercellular transport of the histone-anchor Jabba and thus its presence in the ooplasm. Jabba prevents ooplasmic H2Av from degradation, safeguarding the H2Av stockpile. Our findings provide insight into the mechanism for establishing histone stores during *Drosophila* oogenesis and shed light on the function of LDs as protein-sequestration sites.

## Introduction

Histones perform critical functions in eukaryotic cells, from promoting genome stability (Herrero & Moreno, 2011) to regulating gene expression (Celona et al., 2011). These roles are highly sensitive to histone abundance: both too few and too many histones can result in severe defects, including increased DNA damage sensitivity, enhanced rates of chromosome loss, aberrant nuclear segregation, cell cycle arrest, and altered gene expression (Gunjan & Verreault, 2003; Han et al., 1987; Meekswagner & Hartwell, 1986; Singh et al., 2009). Therefore, in typical somatic cells, histone abundance at any one moment is closely coupled to histone demand. Synthesis of canonical histones, for example, occurs almost exclusively when DNA replication provides a sink for histone deposition, and this S-phase restriction is ensured via multiple regulatory mechanisms that control transcription and mRNA turnover (Marzluff et al., 2008).

However, these powerful regulatory strategies are not available during developmental stages that are transcriptionally silent, such as early embryos. New histones messages cannot be synthesized, and destruction of messages at the end of one S-phase would prevent translation of more histones in the following S-phase. To meet the histone demand of the extremely rapid early embryonic cell cycles, many animal eggs contain large stockpiles of excess canonical and variant histone proteins (O’Farrell et al., 2004). These stockpiles can be impressive; for *Drosophila* embryos, it has been estimated that the newly laid egg has a thousand-fold excess of histones over DNA (Cermelli et al., 2006).

Such stockpiling, however, leads to a new challenge since excess histones are toxic (Gunjan & Verreault, 2003; Han et al., 1987; Marzluff et al., 2008; Meekswagner & Hartwell, 1986; O’Farrell et al., 2004; Singh et al., 2009); therefore, early embryos have to somehow prevent the dangers of histone overabundance.

Two general strategies have been uncovered that organisms employ to solve this problem. Pioneering work in *Xenopus* established that excess histones can be complexed with histone chaperones in the cytoplasm. These complexes are stable, and histone release is developmentally controlled by post-translational modifications of the chaperones (Onikubo et al., 2015). In *Drosophila* embryos, a subset of histones (H2B, H2A, and H2Av) is sequestrated on lipid droplets (LDs) (Cermelli et al., 2006), the cellular organelles specialized for fat storage (Walther& Farese, 2012). LD-bound histones can be transferred to nuclei (Cermelli et al., 2006; Johnson et al., 2018) and contribute to successful early development (Li et al., 2012). Although there has not yet been a comprehensive survey of how widespread either of these two strategies of handling excess histones are, histones on LDs have been detected in embryos and oocytes of house flies and mice (Cermelli et al., 2006; Kan et al., 2012) and even in some somatic cells. For example, H2A and H2B were identified biochemically as bona-fide LD proteins in human osteosarcoma (U2OS) and human hepatocellular carcinoma (Huh7) cell lines (Bersuker et al., 2018), and microscopically in rat sebocytes (Kan et al., 2012; Nagai et al., 2005; for more examples see also Welte, 2015).

LDs have extensively been characterized for their roles in lipid metabolism and energy homeostasis (Walther & Farese, 2012; Yu & Li, 2017; Zechner et al., 2012), but it is increasingly recognized that they also perform important regulatory functions in the life cycles of many proteins (Welte and Gould, 2017). For example, upon Hepatitis C infection, several newly made viral proteins transiently associate with LDs; this step is essential for virion assembly and maturation (Filipe & McLauchlan, 2015). Other proteins, such as excess Apolipoprotein B in hepatocytes, accumulate on LDs before they get degraded (Ohsaki et al., 2006; Ohsaki et al., 2008). It has been proposed that sequestration of ER proteins on LDs and subsequent autophagic degradation of these LDs is a general mechanism to rid the ER of damaged proteins (Vevea et al., 2015). Sequestration of the ER protein Sturkopf to LDs has been proposed to mitigate its repressive effects on an ER-associated enzyme (Werthebach et al., 2019). And a number of proteins involved in transcriptional regulation can be sequestered on LDs in bacteria, fungi, and mammalian cells (Aramburu et al., 2006; Gallardo-Montejano et al., 2016; Mejhert et al., 2020; Romanauska & Kohler, 2018; Zhang et al., 2017). However, in only a few cases is the function of protein sequestration on LDs understood at a molecular level.

In *Drosophila*, sequestration of histones to LDs has two known biological functions, buffering and storage. During oogenesis, LD sequestration somehow boosts overall histone levels, since embryos from mothers lacking Jabba, the histone anchor on LDs, have dramatically reduced levels of H2A, H2B, and H2Av (Li et al., 2012). In early embryos, LD sequestration prevents overaccumulation of newly synthesized histone H2Av on chromatin (Li et al., 2014). The mechanistic basis for buffering has been elucidated: transient binding to LDs retains newly translated H2Av in the cytoplasm, thus slowing its nuclear import (Johnson et al., 2018). However, how H2Av storage is mediated by LDs remains unknown. Addressing this issue could provide an important paradigm for other cases of protein sequestration on LDs.

During *Drosophila* oogenesis, mature eggs are produced by so-called egg chambers that consist of somatic follicle cells and the germ-line derived oocyte and its fifteen sister cells, nurse cells. The oocyte initiates meiosis and is arrested in metaphase (Ables, 2015; Von Stetina & Orr-Weaver, 2011), remaining almost completely transcriptionally silent (Navarro-Costa et al., 2016). It falls to the nurse cells to synthesize a large fraction of the materials and organelles needed for the growth of the oocyte. As nurse cells and the oocyte remain connected through cytoplasmic bridges, nurse cell contents can easily be transferred to the oocyte, initially at a slow steady rate and eventually by the nurse cells dramatically contracting during stages 10b and 11 of oogenesis, which squeezes the remaining nurse cell cytoplasm into the oocyte (Hudson & Cooley, 2014).

This structure of egg chambers suggests three possibilities for how lack of the histone anchor Jabba might lead to reduced histone H2Av levels in newly laid eggs. First, Jabba may be necessary for histone biosynthesis, by directly or indirectly promoting transcription or translation of histones. For example, Jabba might act as transcription co-factor like the LD protein Perilipin 5 that can relocate to the nucleus under conditions of lipolysis (Gallardo-Montejano et al., 2016). As there is some evidence that proteins may be translated at the LD surface (Bozza et al., 2009; Wan et al., 2007), Jabba might, alternatively, recruit histone mRNAs to LDs or regulate their translation. Second, if histones are already made in nurse cells, Jabba may facilitate the transport of histones to the oocyte by attaching them to LDs in the nurse cells. Finally, Jabba may prevent turnover of histones after they have been synthesized, possibly because the surface of LDs is relatively inaccessible to proteases (Keembiyehetty et al., 2011). Indeed, for at least some bona-fide LD proteins, binding to LDs prevents their degradation (Masuda et al., 2006; Rowe et al., 2016; Takahashi et al., 2016; Xu et al., 2006).

Using quantitative imaging, we have determined the developmental time course of H2Av accumulation in oocytes and found that the bulk of oocyte H2Av does not depend on Jabba-mediated transport from nurse cells. We also detect no evidence for Jabba-dependent H2Av biosynthesis but show directly that in the absence of Jabba, H2Av is turned over in a proteasome-dependent manner. Unlike for H2Av, oocyte accumulation of Jabba depends on its ability to bind to LDs. We propose that LDs retain Jabba in the nurse cell cytoplasm, thus allowing its transport into the oocyte, where it then protects H2Av from degradation.

Taken together, our results suggest that during oogenesis LD sequestration is necessary to get the histone partner Jabba into the oocyte, where Jabba then promotes histone stability.

## Results

### Histones localize to nurse cell LDs in a Jabba-dependent manner

We envision that Jabba promotes histone storage by boosting histone synthesis, or promoting histone transport from nurse cells to oocytes, or preventing histone degradation. To start distinguishing between these models, we asked when the association between LDs and histones arises, using females expressing an H2Av-RFP fusion protein and co-labeling fixed samples with the LD-specific dye BODIPY. In agreement with earlier studies (summarized in Welte, 2015) we found that LDs are relatively sparse until stage 8. LDs are present in massive amounts in the nurse cells from stage 9 onwards until dumping, when the nurse cell cytoplasmic contents are transferred into the oocyte (not shown). We similarly observed widespread puncta of H2Av-RFP in the cytoplasm of nurse cells of the same stages (see also Fig. 1.C), and they co-localized with LDs (Fig. 1.A). This result is consistent with earlier observations where LD-localization of H2Av in nurse cells was inferred from centrifugation experiments (Cermelli et al., 2006).

**Figure 1.**
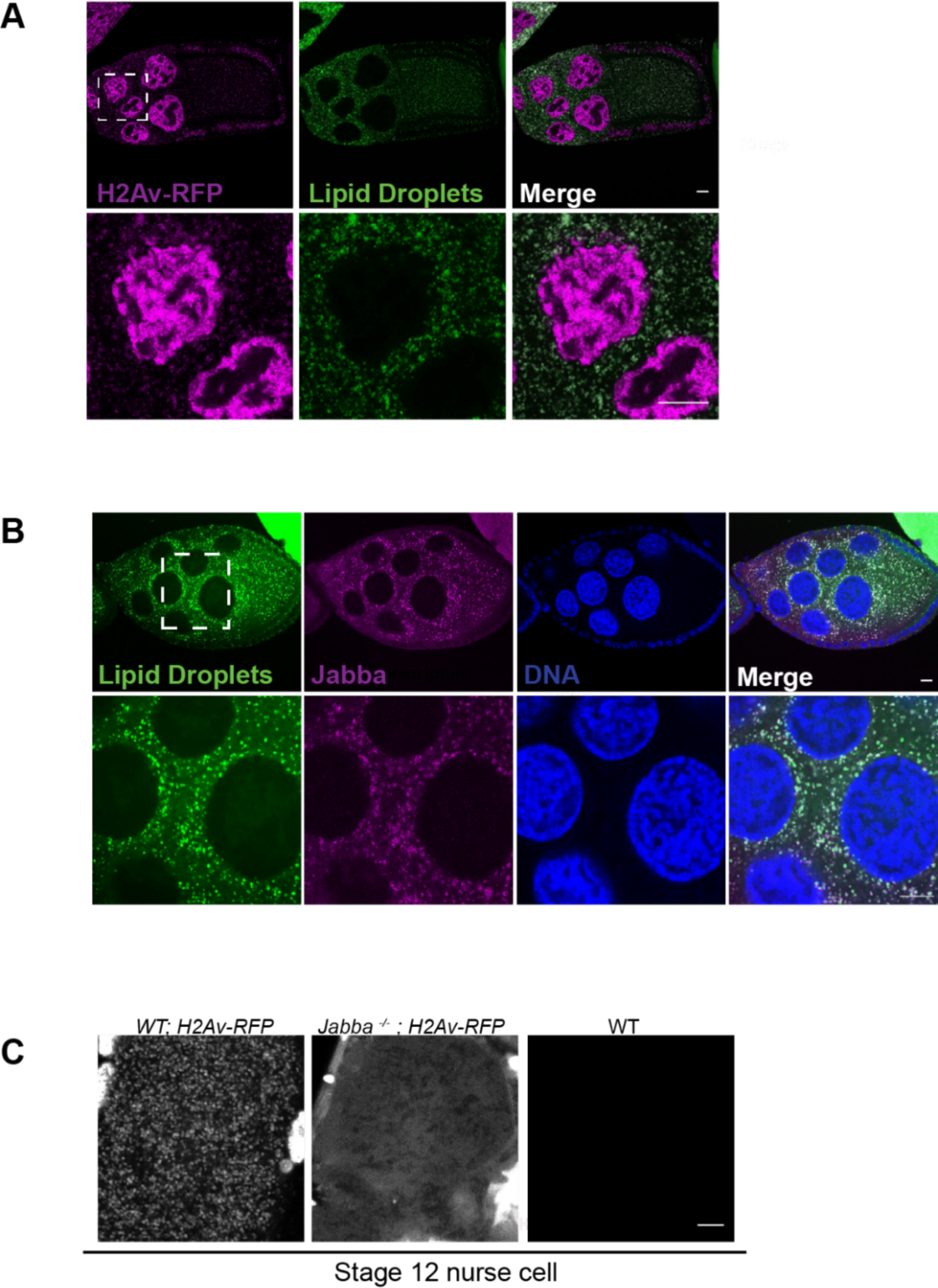
Histones localize to nurse cell LDs in a Jabba-dependent manner. A) H2Av (magenta) is present on lipid droplets (LDs) (green) in the ovary. Stage 9 egg chamber expressing H2Av-RFP was stained with BODIPY to visualize LDs. (B) Jabba (magenta) localizes to LDs (green) as early as LDs accumulate during stage 9. LDs are stained with Nile Red. Nuclei are stained with DAPI (blue). (C) Jabba anchors H2Av to LDs in the ovary. Nurse cells of stage 12 wild-type and *Jabba ^-/-^* egg chambers expressing H2Av-RFP. H2Av fluorescence (white) is present in distinct puncta in the wild type (left) and diffuse through the cytoplasm in *Jabba ^-/-^* (center); wild-type nurse cells not expressing fluorescently tagged H2Av (right) show no signal under identical imaging conditions. Dashed boxes indicate the regions of interest used for higher magnification. Scale bars represent 10 μm.

In the embryo, H2Av-LD association is mediated by Jabba (Li et al., 2012), likely via direct physical interactions (Kolkhof et al., 2017; Li et al., 2012). Immunostaining revealed that Jabba was indeed already present on LDs in nurse cells (Fig. 1.B). By live imaging, H2Av-RFP is present in distinct particles, which represent LDs (Fig. 1.C, left). In nurse cells from *Jabba* mutant egg chambers, no such accumulation was detectable; rather H2Av-RFP signal was diffuse through the cytoplasm (Fig. 1.C, middle). Comparison of a strain not expressing H2Av-RFP (Fig.1.C, right) identifies the diffuse signal as H2Av-RFP rather than background fluorescence. Thus, like in embryos, Jabba is necessary to recruit H2Av to LDs and serves as histone anchor. However, unlike in embryos, where cytoplasmic signal is negligible in the mutants (Johnson et al., 2018; Li et al., 2012), cytoplasmic H2Av is still abundant. These observations suggest that H2Av synthesis is not completely abolished in the absence of Jabba.

### LDs make at most a modest contribution to H2Av accumulation in oocytes

From our data it is clear that LDs and H2Av are produced in nurse cells, but they do not address what fraction of LDs/H2Av present in mature oocytes come from nurse cells. To be able to address the contribution of LD-mediated H2Av transport, we first sought to determine the relationship between the histone pool in the nurse cells and in the oocyte. Using flies expressing H2Av-RFP, we quantitated the relative cytoplasmic concentration of H2Av-RFP at various stages of oogenesis by measuring mean fluorescence intensity in the ooplasm (Fig. 2.A, B). H2Av concentration increased steadily from stage 10a (before dumping, mid-oogenesis) to stage 14 (late oogenesis), despite the fact that total oocyte volume is known to increase substantially between stage 10 to 12 (Jia et al., 2016). After stage 12, oocyte volume is fairly stable (King & Koch, 1963); nevertheless H2Av-RFP concentration increased about 2.5 fold by stage 14 (Fig. 2.A,B). We performed the same experiments using an H2Av-GFP line as well as an H2B-mEos3.2 line and observed similar accumulation patterns (Figs. 2.C, D). Thus, both H2Av and H2B exhibit a major rise in ooplasmic concentration late in oogenesis.

**Figure 2.**
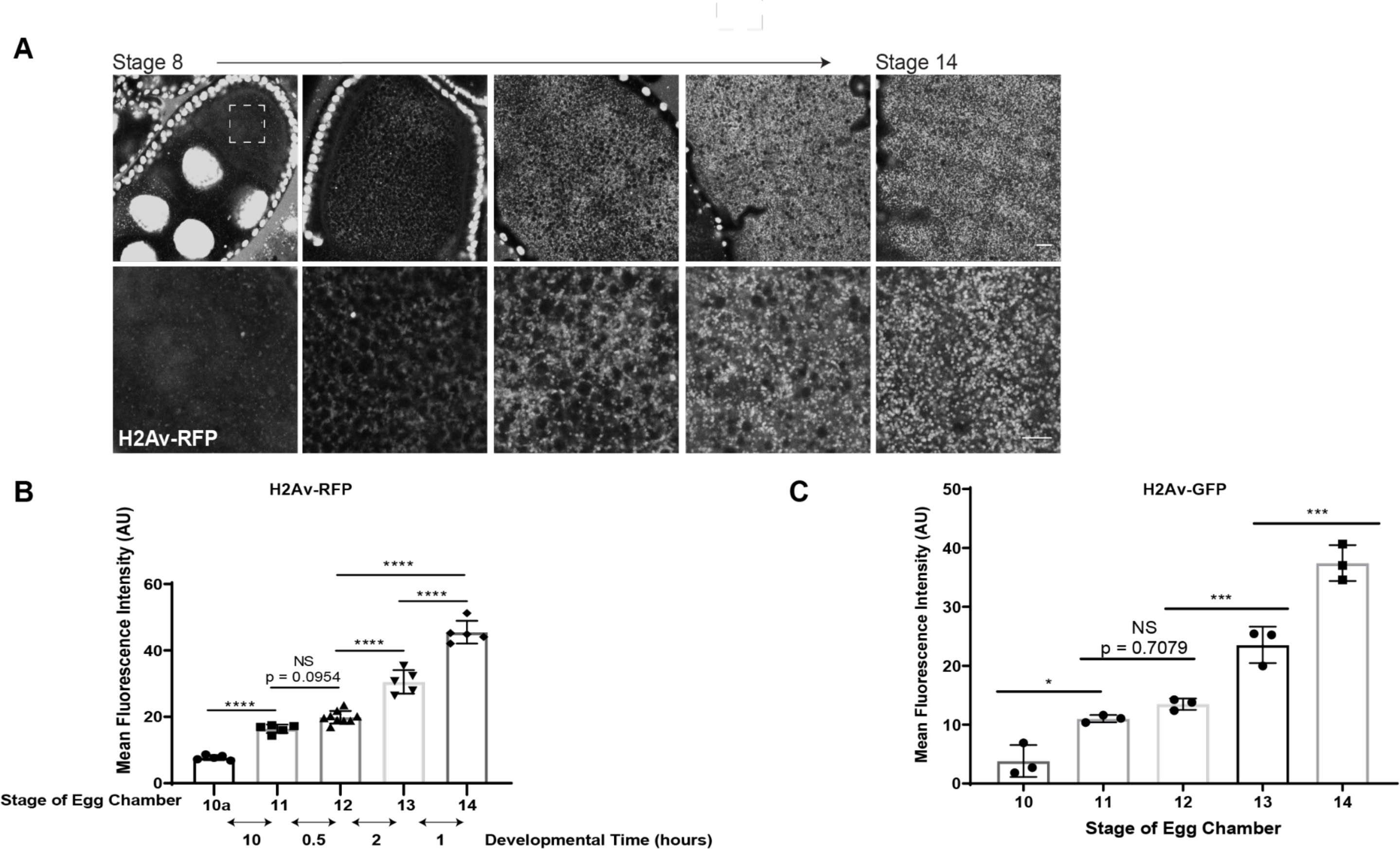
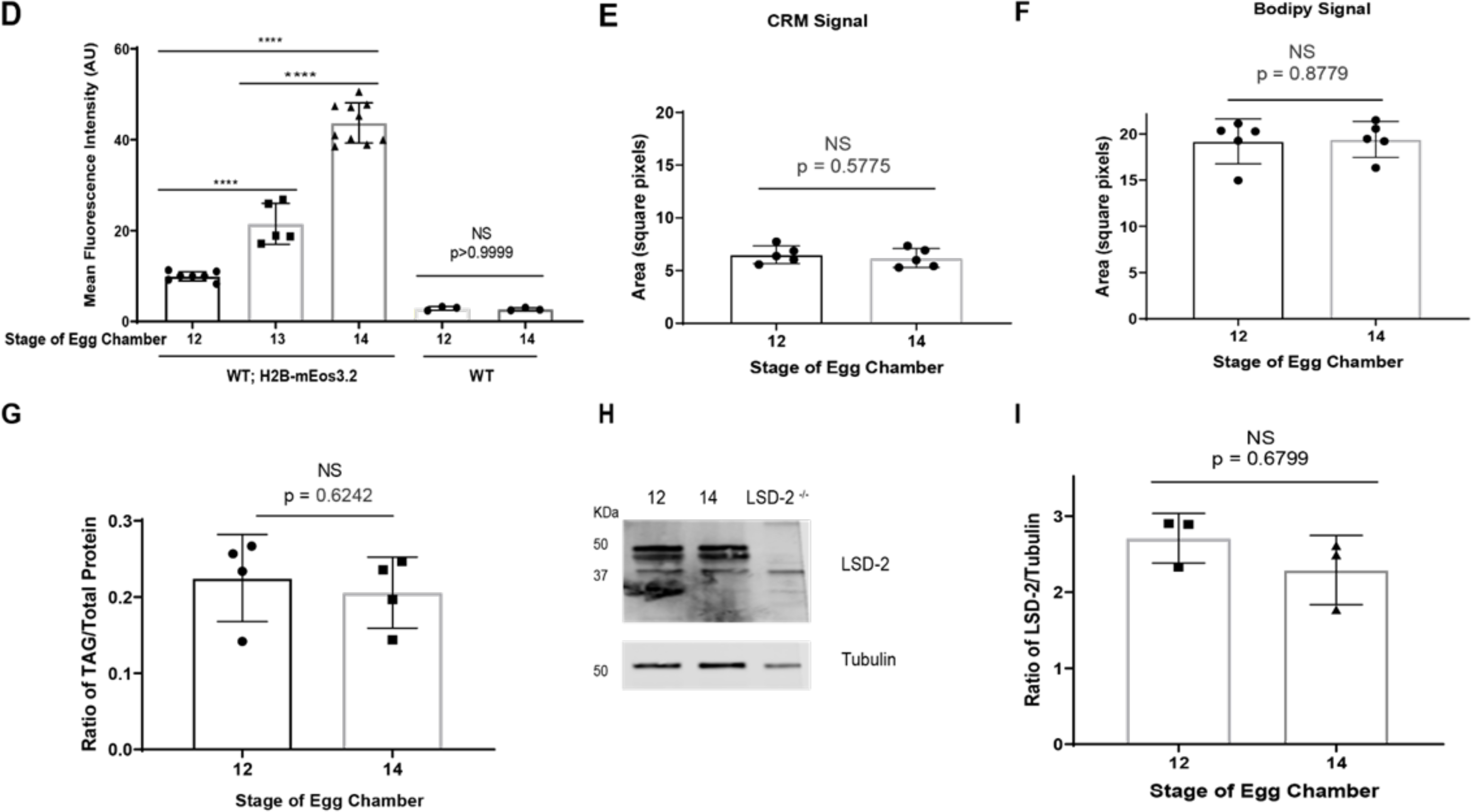
Ooplasmic H2Av levels increase during oogenesis even when LDs remain constant. A-B) H2Av increases over developmental time. Egg chambers of flies expressing H2Av-RFP (white) (A). Dashed box indicates the region of interest in the ooplasm used for higher magnification. Scale bar represents 10 μm. (B) Quantitation of (A) expressed as the Mean Fluorescence Intensity (AU). Double-headed arrows indicate the time of growth (hours) between stages. Error bars represent SD. p values were calculated using a one-way ANOVA followed by Tukey’s test. Stage 10a vs Stage 11, p <0.001; Stage 11 vs Stage 12, p = 0.0954; Stage 12 vs Stage 13, p <0.001; Stage 13 vs Stage 14, p <0.001; Stage 14 vs Stage 12, p <0.001. (C) Quantitation of the Mean Fluorescence Intensity (AU) of H2Av-GFP in the ooplasm over developmental time; the same overall pattern is seen as when H2Av-RFP is employed (in B). p values were calculated using a one-way ANOVA followed by Tukey’s test. stage 10 vs stage 11, p = 0.0243; stage 11 vs stage 12, p = 0.7079; stage 12 vs stage 13, p = 0.0026; stage 13 vs stage 14, p = 0.0002. (D) Quantitation of the Mean Fluorescence Intensity (AU) of H2B-mEos3.2 in the ooplasm over developmental time. Fluorescence Intensity was very low in the WT egg chambers that expressed no fluorescently tagged H2B and those levels did not change significantly between stages. stage 12 vs stage 13, p <0.001; stage 13 vs stage 14, p <0.001; stage 12 vs stage 14, p <0.001; WT stage 12 vs WT stage 14, p > 0.9999. (E-F) The area covered by LDs in the ooplasm of stage 12 and stage 14 egg chambers in images acquired by Confocal Reflection Microscopy (CRM) (E) or after staining with BODIPY (F). (G) The ratio of triglyceride levels relative to total protein in stage 12 and stage 14 egg chambers. (H) Western analysis of LSD-2 levels in stage 12 and stage 14 egg chambers. *LSD-2 ^-/-^* was used as a negative control. Membranes were probed for LSD-2 and for tubulin, as a loading control. (I) Quantitation of (H) expressed as the H2Av/tubulin ratio. N = 3. In E-I, p values were calculated using a Student’s t test.

Although dumping is essentially complete by stage 12, it is theoretically conceivable that some LDs continue to be funneled from nurse cells in the oocyte, contributing little to overall oocyte volume, but bringing in more H2Av. To test this possibility, we quantified LD concentrations in the oocyte cytoplasm from stage 12 to 14. We measured the area covered by LDs in a single focal plane of the ooplasm of stage 12 and stage 14 egg chambers (Fig. 2.E, F). Additionally, we determined the total triglyceride (TAG) levels relative to total protein content in stage 12 versus stage 14 egg chambers (Fig. 2.G). Both measures indicate that the net amount of LDs is unchanged from stage 12 onwards. In addition, the abundance of the LD-bound protein LSD-2, as assessed by Western analysis (Fig. 2.H, I), did not change between stage 12 and stage 14 egg chambers. Collectively, these observations suggest that the majority of the LDs in the oocyte is generated in the nurse cells prior to dumping and that there are no further increases from stage 12 to stage 14.

The dramatic increase in H2Av concentrations from stage 12 to stage 14 together with unchanged LD concentration indicates that H2Av is newly synthesized in the ooplasm and loaded unto pre-existing LDs. Our quantitation suggests that more than half (56%) of the H2Av pool in mature oocytes originates from synthesis in the oocyte itself after dumping is complete.

To test the contribution of LD-mediated transport to the remaining 44% of the H2Av oocyte pool, we compared H2Av-RFP mean fluorescence intensity of wild-type and *Jabba ^-/-^* oocytes in stage 12, *i.e.*, right after dumping. Because in the mutants H2Av is not LD associated, all LD-mediated H2Av transport should be abolished. Nevertheless, the *Jabba ^-/-^* ooplasm displayed considerable H2Av-RFP signal, about 55% of what is found in the wild type (Fig. 3.A, B). Thus, the contribution of Jabba-dependent transport to the final H2Av pool at stage 14 is at most 20% (=45% of 44%), or possibly much less (see discussion).

**Figure 3.**
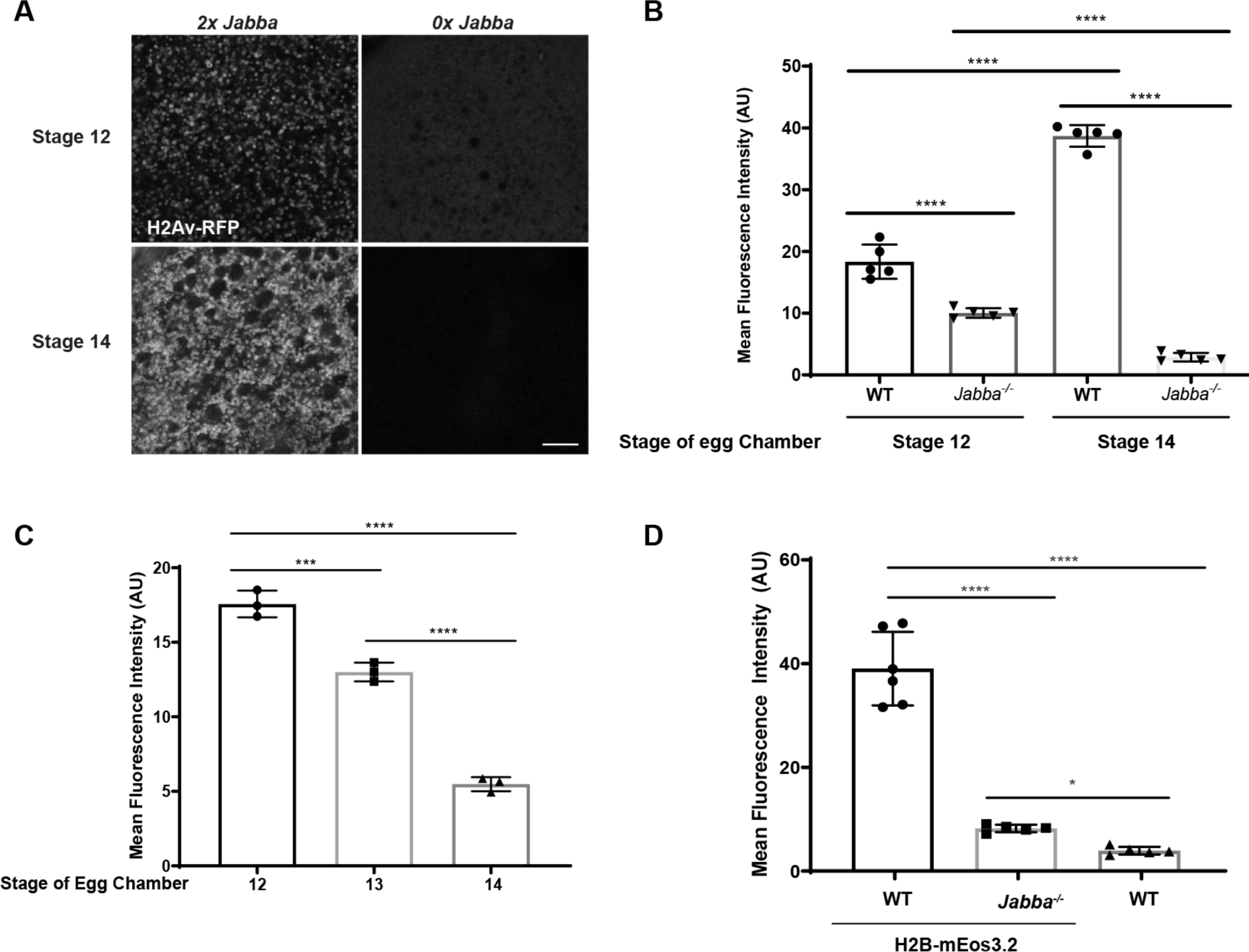
Jabba maintains the maternal supply of H2Av and H2B. A) H2Av-RFP (white) fluorescence in the ooplasm of wild-type and *Jabba ^-/-^* flies during stage 12 and stage 14. Scale bar represents 10 μm. (B) Quantitation of (A) showing Mean Fluorescence Intensity (AU). p values were calculated using a one-way ANOVA followed by Tukey’s test. WT stage 12 vs *Jabba ^-/-^* stage 12, p <0.0001; WT stage 12 vs WT *^stage^* 14, p <0.0001; WT stage 14 vs *Jabba ^-/-^* stage 14, p <0.0001; Jabba *^-/-^* stage 12 vs *Jabba ^-/-^* stage 14, p <0.0001. (C) Mean Fluorescence Intensity (AU) of H2Av-RFP in *Jabba ^-/-^* egg chambers from stage 12 to stage 14. p values were calculated using a one-way ANOVA followed by Tukey’s test. stage 12 vs stage 13, p = 0.0005; stage 12 vs stage 14, p <0.001; stage 13 vs stage 14, p < 0.001 (D) Quantitation of the Mean Fluorescence Intensity (AU) of H2B-mEos3.2 in the ooplasm of stage 14 wild-type and *Jabba ^-/-^* egg chambers. WT egg chambers lacking fluorescently tagged H2B were imaged and analyzed to determine the background signal. p values were calculated using a one-way ANOVA followed by Tukey’s test*. WT; H2B-*mEos3.2 vs WT, p <0.0001; *WT; H2B-mEos3.2* vs *Jabba ^-/-^; H2B-mEos3.2,* p <0.0001; *Jabba ^-/-^; H2B-mEos3.2* vs WT, p= 0.0149

### Jabba prevents the loss of H2Av as oocytes mature

Previous Western analysis had found that in the absence of *Jabba* newly laid eggs have dramatically reduced total amounts of H2Av (Li et al., 2012). Here we find, in contrast, that lack of *Jabba* reduces H2Av concentration in stage 12 oocytes only modestly and that this reduction represents only a small fraction of the total H2Av in mature oocytes. This disparity in results could be due to a technical difference in how we detect H2Av (Western versus imaging) or because H2Av levels in the two genotypes diverge between stage 12 and egg laying. Western analysis of stage 12 oocytes is challenging as it would require removing the degenerating nurse cell nuclei; their high histone levels due to massive polyploidy (King, 1970) would likely overwhelm the oocyte histone signal, even if there is only mild contamination. We therefore opted to assess the developmental time course of H2Av levels by quantitative imaging of the ooplasm.

As described before (Fig. 2.A, B), H2Av-RFP signal in wild-type oocytes goes up from stage 12 to 14 (Fig. 3.A, left). When *Jabba ^-/-^* oocytes are imaged under the same conditions, stage 12 oocytes still show substantial signal, though it is broadly diffuse in the ooplasm instead of being concentrated in distinct puncta. Thus, like in nurse cells, Jabba is apparently required to recruit H2Av to LDs. In stage 14, however, *Jabba ^-/-^* had drastically lower H2Av levels (Fig. 3.A, right). Quantitation revealed that levels had dropped to 28% of the *Jabba ^-/-^* stage 12 levels and to 7% of the wild-type stage 14 levels (Fig. 3.B) and that the drop was gradual, with stage 13 displaying intermediate levels (Fig. 3.C). Thus, in the absence of Jabba, H2Av stores in the oocyte are not stably maintained, and this loss even counteracts the rise usually observed in the wild type. We observed a similar pattern using a tagged H2B construct (Fig. 3.D). Thus, we conclude that the reduction in the maternal pool of histones observed in *Jabba ^-/-^* embryos is already established during oogenesis.

In principle, the divergence in H2Av levels between wild type and *Jabba ^-/-^* could be due to decreased H2Av synthesis or enhanced H2Av degradation in the mutant, or a combination of both. However, there is no evidence that lack of Jabba alters transcription of histone messages or their stability: by qPCR, the levels of H2A, H2AB, and H2Av messages in stage 14 were indistinguishable between wild-type and *Jabba ^-/-^* egg chambers (Figs. 4.B, B’, B’’) or in unfertilized embryos of these genotypes (Figs. 4.A, A’, A”; see also (Li et al., 2012). In addition, immunostaining reveals no appreciable presence of Jabba in the nucleus, as would be expected if it were directly involved in transcription regulation (Fig. 1.B).

To determine whether the reduced histone pool in *Jabba ^-/-^* arises due to impaired translation, we examined the histone mRNAs bound to polysomes. There was no significant change in the H2A, H2Av, and H2B mRNA levels being actively translated in the wild-type and *Jabba ^-/-^* stage 14 egg chambers (Fig. 4.C - C’’). Therefore, we have no evidence that histones synthesis in the absence of Jabba is appreciably altered.

**Figure 4.**
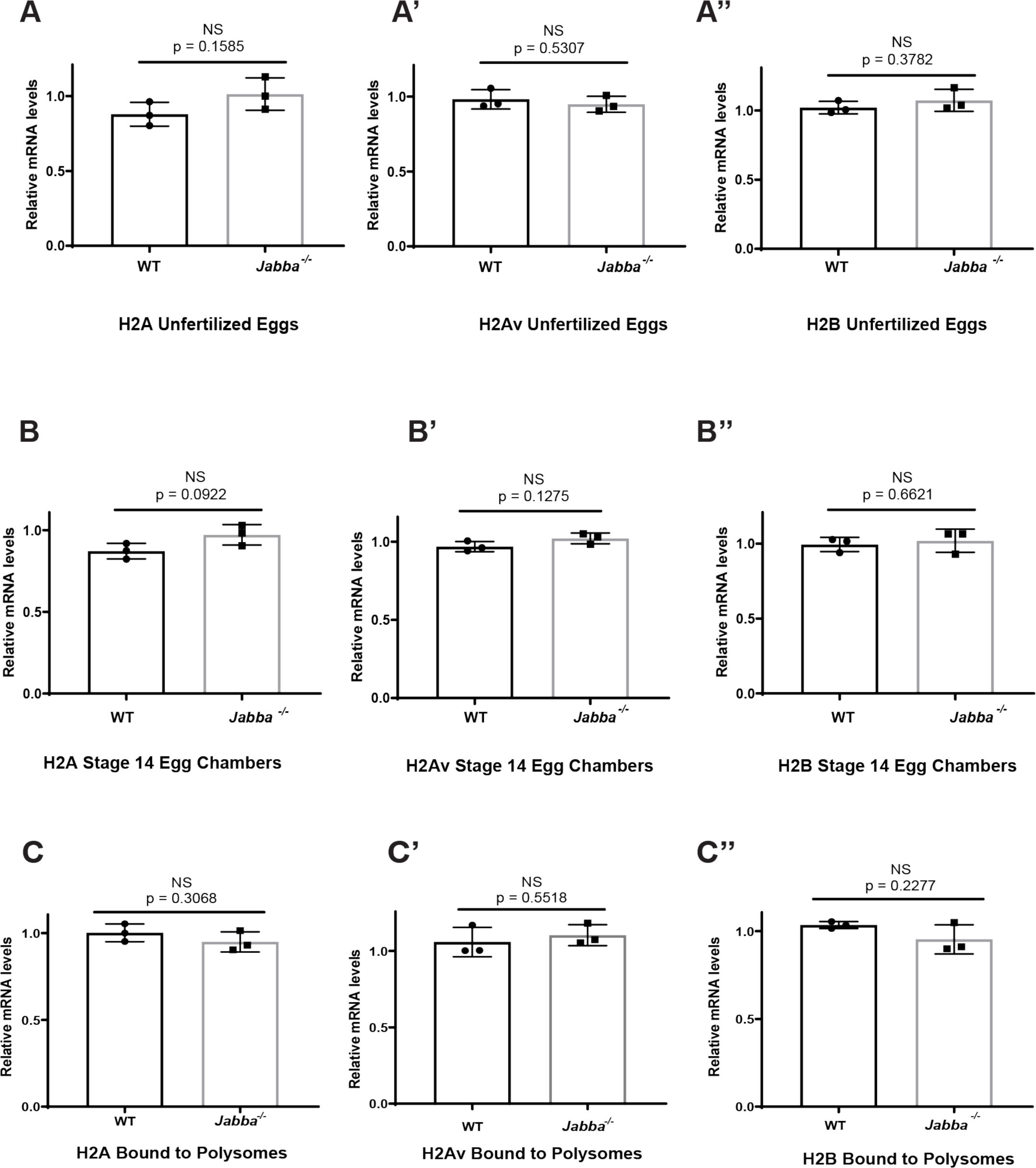
H2A, H2Av, and H2B biosynthesis appears unaffected in *Jabba ^-/-^*. Relative mRNA levels of H2A, H2Av, and H2B in wild-type and *Jabba ^-/-^* unfertilized eggs (A), stage 14 egg chambers (B) and bound to polysomes (C). There is no significant change in relative mRNA levels for any of these histones between wild-type and *Jabba ^-/-^* under the three conditions. p values were calculated using a Student’s t test. Error bars represent SD.

### H2Av is degraded in the absence of Jabba

Our analysis so far indicates that the Jabba-dependence of histone levels in mature oocytes is not due to major roles of Jabba in histone synthesis or transport. These observations suggest that Jabba somehow counteracts the turnover of these histones. If correct, inhibition of histone degradation should be able to reverse the effects of lack of Jabba on histone levels. Since in many systems excess histones are turned over by proteasome-mediated degradation (Imschenetzky et al., 1997; Lin et al., 1991; Morin et al., 2012; Singh et al., 2009), this pathway is an obvious candidate for mediating histone degradation in oocytes. We therefore opted to pharmacologically inhibit the proteasome using *in vitro* egg chamber maturation (IVEM) (Spracklen & Tootle, 2013), to avoid secondary effects on whole-organism viability.

We dissected ovaries from H2Av-RFP expressing females and isolated stage 12 egg chambers (Fig. 5.A). These egg chambers were then incubated in IVEM media and placed in the dark to allow for *in vitro* development. Under these conditions, egg chambers fully progress to stage 14 in a few hours. To determine H2Av-RFP levels, we measured mean ooplasmic fluorescence either in newly dissected egg chambers or after six hours, when full maturation had occurred. The disparity of H2Av-RFP signal between stage 14 wild-type and *Jabba ^-/-^* egg chambers observed *in vivo* was also present under these *in vitro* culture conditions (Fig. 5.B, condition labeled “untreated”), even when the solvent Dimethyl sulfoxide (DMSO) was added to the culture media (Fig. 5.B, condition labeled “0 µg/ml”). These observations set the stage to test the effects of various concentrations of the proteasome inhibitor MG132 (dissolved in DMSO).

**Figure 5.**
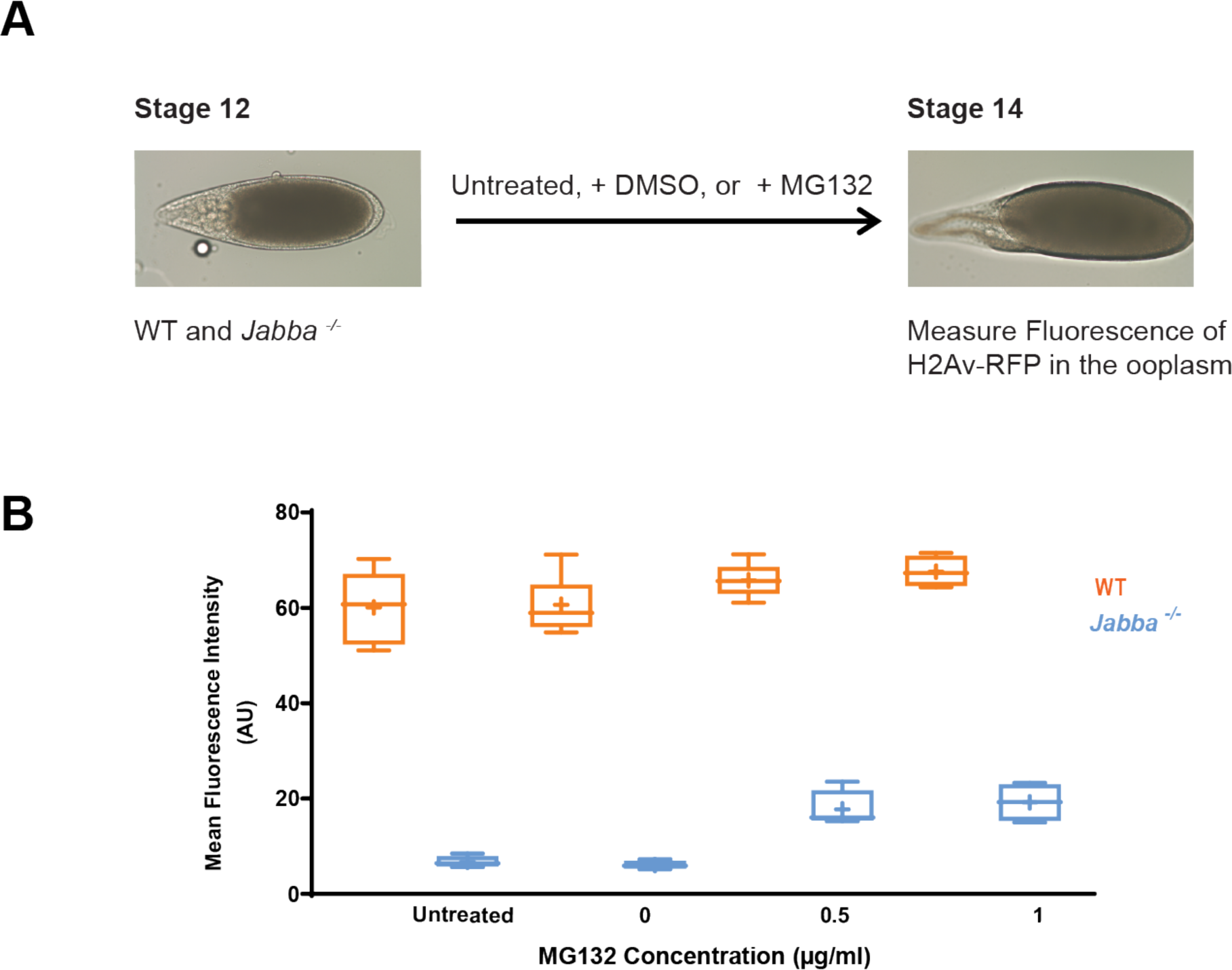
H2Av is degraded in the absence of Jabba. A) Schematic depiction of *in vitro* egg maturation (IVEM) experiment. (B) The mean fluorescence intensity (AU) in stage 14 WT (orange) or *Jabba ^-/-^* (blue) egg chambers after IVEM. Error bars represent SD. Length of box plot represents the 25^th^ and 75^th^ percentile. Line (in box plot) represents median, Cross indicates the mean. Untreated = *in vitro* culture in IVEM media; 0, 0.5, 1 µg/ml= indicates the various concentrations of MG132 (diluted in DMSO) used in IVEM media.

Full inhibition of the proteasome is known to result in developmental arrest during oogenesis (Velentzas et al., 2011). It is therefore not surprising that in our assay high levels of MG132 (50 µg/ml) interfered with egg chamber maturation and egg chambers never reached stage 14. Consistent with arrest, H2Av-RFP levels were comparable to stage 12 egg chambers (untreated) for both wild type and *Jabba ^-/-^* (Fig. 5 - Sup. Fig. 1.A).

We therefore tittered MG132 concentrations down to a point where development to stage 14 was still supported (0.5 and 1 µg/ml). Presumably, under these conditions proteasome function is only partially abolished. H2Av-RFP levels in wild-type egg chambers were largely unaltered. In contrast, H2Av-RFP intensity was decreased in the *Jabba ^-/-^* egg chambers. This difference between genotypes is particularly obvious when we compute the ratio of H2Av-RFP signal in mutant versus wild-type ooplasm (Fig. 5 - Sup. Fig. 1.B). Over a range of inhibitor concentrations, this ratio remains close to the stage 12 and higher than the stage 14 levels, whether the inhibitor still allowed morphological maturation (0.5 and 1 µg/ml) or arrested it (2.5, 5, and 25 µg/ml). We conclude that in the absence of Jabba, H2Av is particularly prone to turnover, consistent with the model that Jabba protects H2Av from degradation.

### H2Av levels in stage 14 oocytes scale with Jabba protein levels

How might Jabba prevent histone degradation? It might act by triggering a molecular switch that regulates the degradation machinery or by physically protecting histones from degradation. To distinguish between these models, we tested whether protection by Jabba scales with the amount of Jabba present and whether it depends on Jabba’s ability to physically bind histones.

If Jabba directly protects histones from degradation, then the level of H2Av in stage 14 oocytes should strongly depend on how much Jabba protein is present; if Jabba acts as a regulatory switch, no such dependency is expected. Jabba protein levels have previously been shown to scale with *Jabba* gene dosage in early embryos (Johnson et al., 2018), and we find the same dependency for oocytes (Fig. 6). When we measured levels of H2Av-RFP by quantitative imaging, we found that mature oocytes from mothers with a single copy of *Jabba* have H2Av levels about half-way between those lacking Jabba entirely and those with two gene copies (Fig. 6.A, B). Remarkably, H2Av levels can even be elevated beyond wild-type levels simply by increasing Jabba expression. As assessed by Western analysis, stage 14 egg chambers from mothers with two additional copies of *Jabba* (*4x Jabba*) accumulate not only more Jabba protein (Fig 7 - Sup. Fig. 1.C, D) but also about twice as much H2Av as the wild type (Fig. 6.C, D). Jabba is limiting for histone accumulation. In fact, increased *Jabba* dosage is even more effective at increasing total H2Av levels than increased *H2Av* dosage (Fig. 6.E); the fact that *H2Av* dosage has any effect on H2Av levels may be due to extra H2Av displacing H2A from their common binding site on Jabba (Kolkhof et al., 2017). Our data is consistent with previous findings which demonstrate that varying *Jabba* dosage in the embryo modulates how much histone can be bound by LDs (Johnson et al., 2018). Taken together, these results suggest that Jabba directly protects H2Av from degradation.

**Figure 6.**
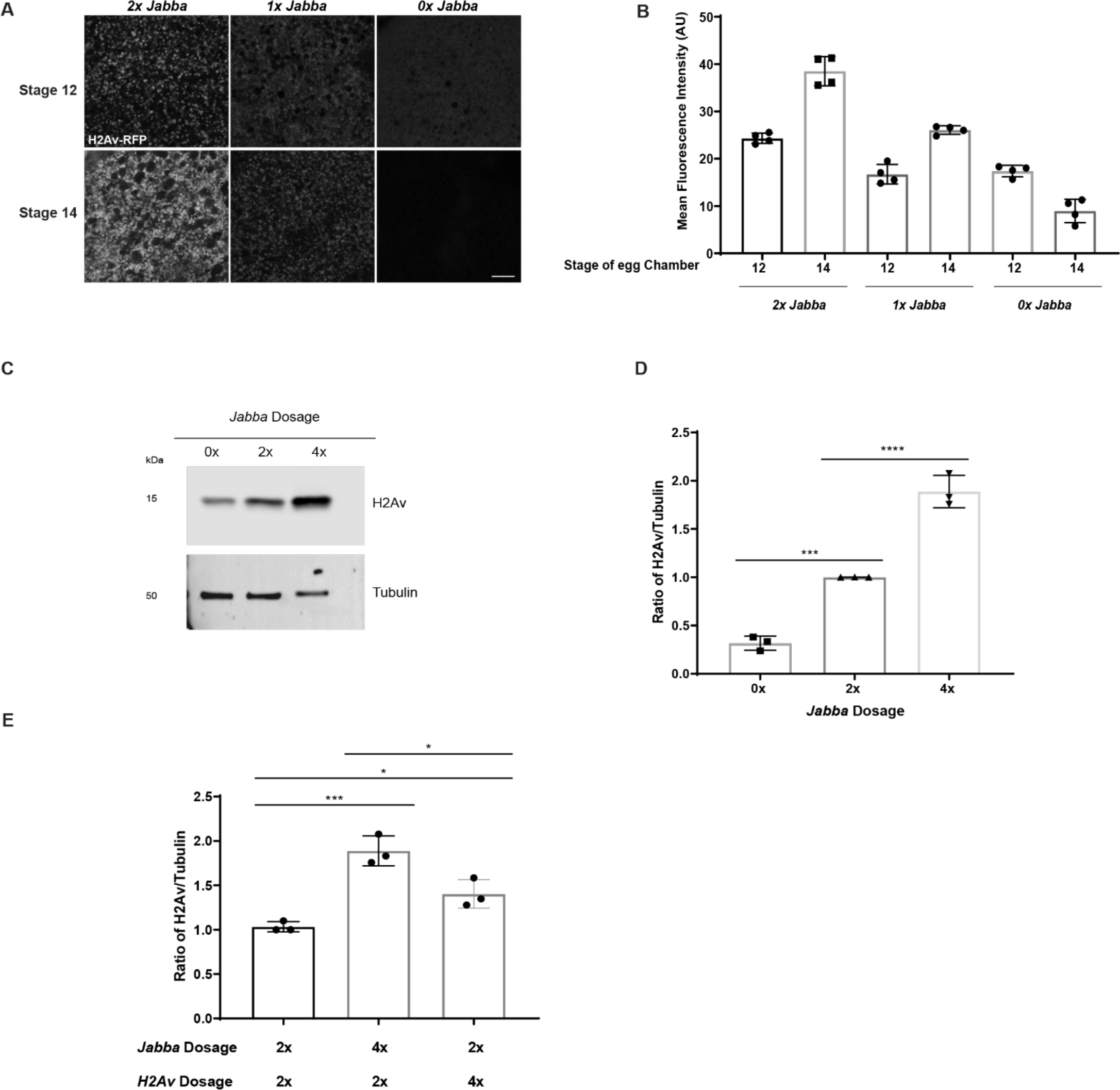
H2Av supply scales with *Jabba* dosage. A) H2Av-RFP (white) fluorescence in the ooplasm of egg chambers expressing 0,1 or 2 copies of *Jabba* during stage 12 and stage 14. Scale bar represents 10 μm. (B) Quantitation of (A) showing mean fluorescence intensity (AU). A subset of these data is presented in Fig. 3.A, B. (C) Western analysis of H2Av levels in stage 14 egg chambers of flies expressing 0, 2 or 4 copies of *Jabba*. Membranes were probed for H2Av and for tubulin, as a loading control. (D) Quantitation of (C) expressed as the H2Av/tubulin ratio. N = 3. p values were calculated using one-way ANOVA followed by Tukey’s test. *0x Jabba* vs *2x Jabba,* p = 0.0002; *2x Jabba* vs *4x Jabba,* p < 0.0001.(E) Quantitation of anti-H2Av Western analysis for the indicated genotypes. Total H2Av is expressed as the ratio of H2Av/tubulin. N = 3. p values were calculated using one-way ANOVA followed by Tukey’s test. *2x Jabba;2x H2Av* vs *4x Jabba;2x H2Av,* p = 0.0007; *2x Jabba;2x H2Av* vs *2x Jabba;4x H2Av,* p = 0.0388; *4x Jabba;2x H2Av* vs *2x Jabba;4x H2Av,* p = 0.012.

**Figure 7.**
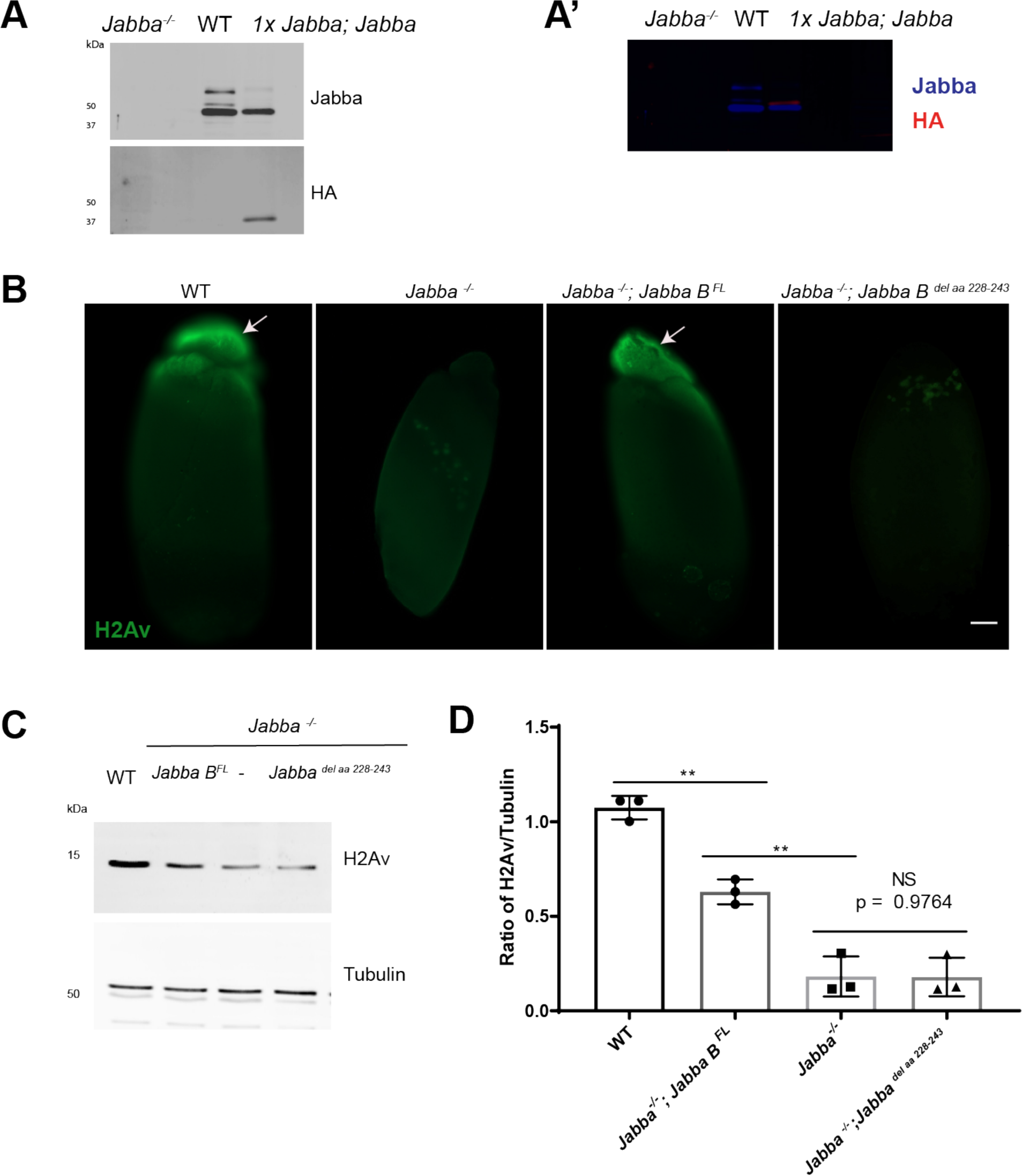
Jabba protects H2Av from degradation. A) Lysates from embryos from *Jabba ^-/-^*, wild-type, and *1xJabba; Jabba B-FLAG-HA* mothers were probed with anti-Jabba (top) and anti-HA (bottom). In the wild-type, three major Jabba bands are visible. Right panel (A’) is an overlay of the anti-Jabba (blue)and anti-HA signal (red). This demonstrates that Jabba B-FLAG-HA comigrates with the lowest of the three endogenous Jabba bands. Based on the apparent molecular weight of Jabba B, the apparent molecular weights of the top bands are consistent with them representing Jabba H and Jabba G. (B) Anti-H2Av immunostaining of centrifuged embryos. H2Av is present in the LD layer (arrow) in WT and embryos expressing *Jabba ^-/-^; Jabba B ^FL^.* Histones are absent from the LD layer in *Jabba ^-/-^* and *Jabba ^-/-^; Jabba B ^del aa 228-243^* embryos. Scale bar represents 50 μm. (C) Western analysis of H2Av levels in stage 14 egg chambers of WT, *Jabba ^-/-^*, *Jabba ^-/-^; Jabba B ^FL^* and *Jabba ^-/-^; Jabba B ^delaa 228-243^* flies. Membranes were probed for H2Av and for tubulin, as a loading control. (D) Quantitation of (C) expressed as the H2Av/tubulin ratio. N = 3. p values were calculated using one-way ANOVA followed by Tukey’s test. *WT* vs *Jabba ^-/-^; Jabba B ^FL^,* p = 0.0002; *WT* vs *Jabba ^-/-^*, p <0.0001; WT vs *Jabba ^-/-^;Jabba ^del aa228-243^,* p <0.0001; *Jabba ^-/-^; Jabba B* ^FL^ vs *Jabba^-/-^*, p = 0.0124; *Jabba ^-/-^* ;*Jabba B ^FL^ vs Jabba ^-/-^;Jabba ^del aa228-243^*, p = 0.0118; *Jabba ^-/-^ vs Jabba ^-/-^;Jabba ^del aa228-243^,* p >0.9999

### H2Av binding is necessary for Jabba’s protective effect

Using a Split Luciferase Protein-Protein Interaction Assay, we had previously shown in *Drosophila* Kc167 cells that Jabba-H2Av interactions require a short, positively charged stretch of amino acids in Jabba (Kolkhof et al., 2017). We therefore set out to develop a platform to determine if the same mutation abolishes H2Av interactions *in vivo* and then test how that affects H2Av accumulation in oogenesis.

We first developed a simplified platform to analyze the ability of mutant Jabba constructs to localize to LDs and recruit histones. The endogenous Jabba locus gives rise to 8 different messages that are predicted generate 7 distinct proteins (Li et al., 2012). These protein isoforms all share an N-terminal region of 316 amino acids and differ in their C-terminal tails. Publicly available RNA-seq data from Flybase (Thurmond et al., 2019) of late egg chambers and early embryos indicate that the predominant isoforms are Jabba B, Jabba G, and Jabba H, with Jabba B most abundant. Consistent with this notion, Western analysis of these stages reveals three major Jabba bands, and the most abundant one co-migrates with ectopically expressed HA-tagged Jabba B (Fig. 7.A).

Using the Gal4/UAS system, we next expressed an mCherry tagged full length Jabba B (Jabba B ^FL^) fusion protein in an otherwise *Jabba* null genetic background and analyzed its ability to bind to LDs using *in vivo* embryo centrifugation. In this assay, cellular components separate by their densities within an intact embryo and LDs are enriched at the anterior end of the embryo (Tran & Welte, 2010). Jabba B ^FL^ prominently accumulated in this lipid layer (Fig. 7 - Sup. Fig. 2.A), demonstrating that it is associated with LDs. Immunostaining of such embryos also revealed that the construct rescued H2Av recruitment to LDs (Fig. 7.B). Using an imaging approach, we compared the expression level of Jabba B ^FL^ to that of endogenous Jabba and found that is comparable to that from a single copy of Jabba (1x Jabba, Fig. 7 - Sup. Fig. 1.D-H). Finally, we measured H2Av levels in stage 14 egg chambers by Western analysis and found that Jabba B ^FL^ was able to maintain H2Av to a similar level as *1x Jabba* (Fig. 7C, D). These observations suggest that Jabba B can protect histones from degradation in the absence of other Jabba isoforms and give us a platform to introduce mutations into Jabba proteins and test their effect on histone metabolism.

We next generated a similar transgene, but with the positive stretch of amino acids (aa) deleted, *Jabba B ^del aa 228-243^.* The mutant protein was expressed at similar levels as Jabba B (Fig. 7 - Sup. Fig. 1.G,H) and localized to LDs in the *in vitro* centrifugation assay (Fig. 7 - Sup. Fig. 2.A). However, it failed to restore H2Av recruitment to LDs (Fig. 7.B), suggesting that aa 228-243 are also necessary for histone binding *in vivo.* In addition, H2Av protection in oogenesis was not rescued; the H2Av pool in *Jabba B ^del aa 228-243^* was similar to that of *Jabba ^-/-^* egg chambers (Fig. 7.C, D). Thus, it is unlikely that Jabba acts as a switch for maintaining histones; rather, we propose that Jabba physically protects H2Av from degradation.

### LD binding allows Jabba to be transported into the oocyte

If Jabba’s primary role is indeed to protect histones from degradation, it is not clear why Jabba is present specifically on LDs. In principle, Jabba free in the ooplasm/not bound to LDs should be sufficient to protect the histone pool from degradation. We therefore designed a Jabba truncation that lacks the previously identified motif for LD recruitment, but retains the stretch of positively charged amino acids important for histone binding. When analyzed in Kc167 cells, this fragment (Jabba[aa192-320]) indeed is not localized to LDs, but present broadly throughout the cytoplasm and the nucleus (Fig. 8.A). By luciferase complementation assay, it robustly interacts with H2Av (Fig. 8.B, Fig 8 - Sup. Fig. 1.A). We will therefore refer to this fragment in the following as Jabba ^HBR^, for histone binding region.

**Figure 8.**
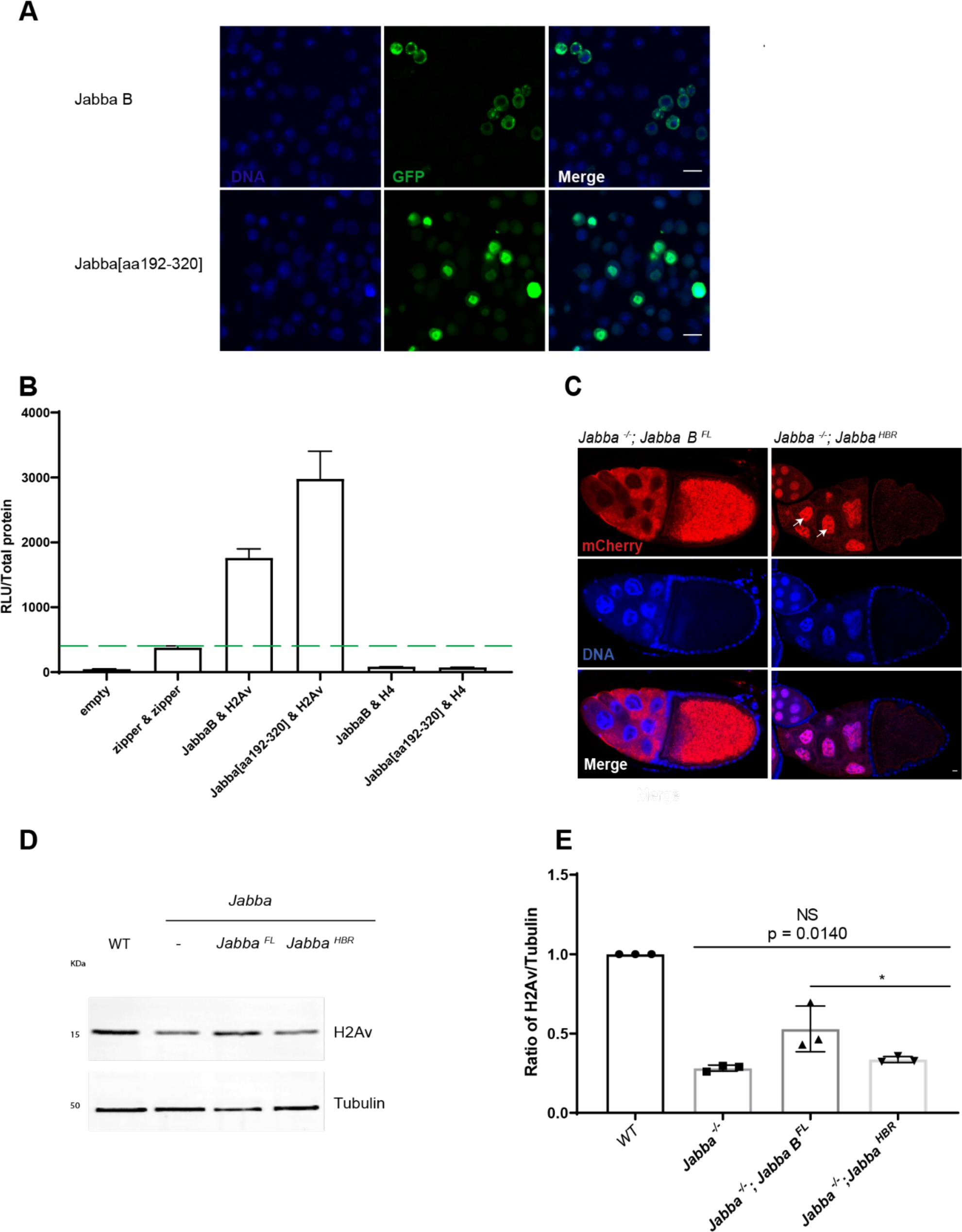
LD binding allows Jabba to be transported into the oocyte. A) Localization of Jabba [aa192-320] in *Drosophila* Kc167 tissue culture cells. Jabba B (green, top panel) is present in the cytoplasm. Jabba [aa192-320] (green, bottom panel) is enriched in nuclei (blue). Scale bar represents 10 μm. (B) Split luciferase complementation assay showing results for the co-expression of the indicated proteins. Luciferase complementation readings are expressed as relative light units (RLU) per μg total protein (RLU/Total protein). Error bars indicate the standard error of the mean (SEM). The green dashed line represents the average of two “zipper-zipper” controls (only one of which is shown), which was used as a threshold for positive interactions. (C) DAPI staining (blue) of egg chambers expressing mCherry tagged Jabba B ^FL^ and Jabba ^HBR^ in a *Jabba ^-/-^* genetic background. mCherry signal (red) is enriched in nurse cell cytoplasm and ooplasm in *Jabba ^-/-^; Jabba B ^FL^*. In *Jabba ^-/-^; Jabba ^HBR^* egg chambers, mCherry signal is present in nurse cell nuclei (arrows). (D) Western analysis of H2Av levels in stage 14 egg chambers of WT, *Jabba ^-/-^, Jabba ^-/-^; Jabba B ^FL^* and *Jabba^-/-^; Jabba ^HBR^* flies. Membranes were probed for H2Av and for tubulin, as a loading control. (E) Quantitation of (D) expressed as the H2Av/tubulin ratio. N = 3. p values were calculated using one-way ANOVA followed by Tukey’s test. *Jabba ^-/-^* vs *Jabba ^-/-^; Jabba ^HBR^*, p = 0.0140; *Jabba ^-/-^; Jabba B ^FL^* vs *Jabba ^-/-^; Jabba ^HBR^*, p = 0.0486

Jabba B ^FL^ expressed during oogenesis (in an otherwise *Jabba ^-/-^* background) mimicked the distribution of endogenous Jabba: Jabba B ^FL^ was detected in the nurse cell cytoplasm and the ooplasm. In contrast, Jabba ^HBR^ was highly enriched in the nurse cell nuclei, with a small amount present diffuse in the nurse cell cytoplasm. By imaging, we detected only minor amounts of Jabba ^HBR^ in the oocyte nucleus (Fig. 8.C).

Although we do not know yet why Jabba ^HBR^ accumulates in nuclei, we speculate that it may reflect in part its ability to bind to histones. Our observations suggest that the reason Jabba is localized to LDs is to prevent it from becoming trapped in nurse cell nuclei which would prevent its transport into the oocyte.

The misdistribution of Jabba ^HBR^ presented us with the opportunity to test whether Jabba has to be physically present in the oocyte to protect histones from degradation. The amount of H2Av detected in stage 14 *Jabba ^HBR^* oocytes was comparable to that for *Jabba ^-/-^* and lower than for *Jabba B ^FL^* (Fig. 8.D, E). Thus, a version of Jabba able to bind to H2Av, but not present in the oocyte, is unable to prevent H2Av degradation. This conclusion is consistent with the notion that Jabba physically protects H2Av in the oocyte cytoplasm.

## Discussion

### Histone stores accumulate from mid to late oogenesis

Early animal embryogenesis is often extremely quick, with very rapid cycles dominated by DNA replication and mitosis and little to no transcription.

*Drosophila* is a particular dramatic case where the first 13 nuclear divisions occur near simultaneously every 8-20 minutes (Foe & Alberts, 1983; Schejter & Wieschaus, 1993). This streamlined development poses a unique problem for histone biology: histone demands goes up exponentially, doubling with each division, yet major regulatory mechanisms that control histone expression in somatic cells are unavailable because they rely on controlling histone mRNA levels. To meet this demand, *Drosophila* embryos – like those of many other species – inherit large, maternally synthesized histone stockpiles. Here, we examined the origin of the maternal stockpile for the variant histone H2Av as well as that for the classical histones H2B and H2A, those stockpiles that depend on the LD protein Jabba (Li et al., 2012).

In particular, we examined when during oogenesis these stockpiles are set up. It has long been known that expression of histone messages is massively ramped up during stage 10 of oogenesis (Ambrosio & Schedl, 1985; Ruddell & Jacobs-Lorena, 1985), likely the source of the maternally provided histone messages in the early embryo. However, it was unclear exactly when the complementary pool of maternally deposited histone proteins accumulates. One of the methodological challenges is that egg chambers have polyploid nuclei. Each of the fifteen nurse cell nuclei have been estimated to contain about 500 times more DNA than diploid nuclei (King, 1970); the 900 follicle cells (Fadiga & Nystul, 2019) contain DNA levels ranging from 8-16 times more DNA than a diploid nucleus (Mulligan & Rasch, 1985). Thus the histones needed to package the nurse cell and follicle cell chromatin are likely at least tenfold more abundant than the histone stockpile in the new laid embryo, which has been estimated to be the equivalent of on the order of 1000 diploid nuclei (Cermelli et al., 2006). Detecting accumulation of the oocyte histone stockpile on top the other histones present in the egg chamber is therefore challenging.

We followed the accumulation of the H2Av stockpile using an imaging approach that specifically quantifies histone signal in the cytoplasm of nurse cells and oocytes. We found that H2Av already accumulates in the cytoplasm of stage 9 nurse cells, well before nurse cell dumping, and that the cytoplasmic H2Av is associated with LDs. This H2Av-LD association likely brings some H2Av into the oocytes; indeed, live imaging suggests that H2Av-RFP loaded LDs can pass through ring canals from nurse cells to oocytes (Jonathon Thomalla and Michael Welte, unpublished observations). However, it is conceivable that almost all of the H2Av present in oocytes is synthesized there; in this case, the reduced H2Av levels in *Jabba ^-/-^* stage 12 oocytes (Fig. 3.A, B) would represent early loss by degradation rather than aberrant H2Av accumulation in nurse cell nuclei or defective transfer from nurse cells. In any case, our quantitative analysis indicates that transfer from nurse cells contributes at most a fifth of the final H2Av pool present in mature oocytes.

After dumping, total H2Av levels continue to rise from stage 12 through stage 14. We propose that two different mechanisms contribute to this rise. First, previous work has found that the translational efficiency of H2Av mRNA is massively upregulated from stage 12 to stage 14 (Eichhorn et al., 2016). Second, Jabba levels increase from stage 12 to stage 14 (Fig 7 - Sup. Fig. 1.A, B). As more Jabba protein becomes available, it can presumably recruit more H2Av to LDs and protect it from degradation (Fig. 9.A’). Our dosage studies indeed indicate that *Jabba* is limiting for H2Av accumulation (Fig. 6).

**Figure 9.**
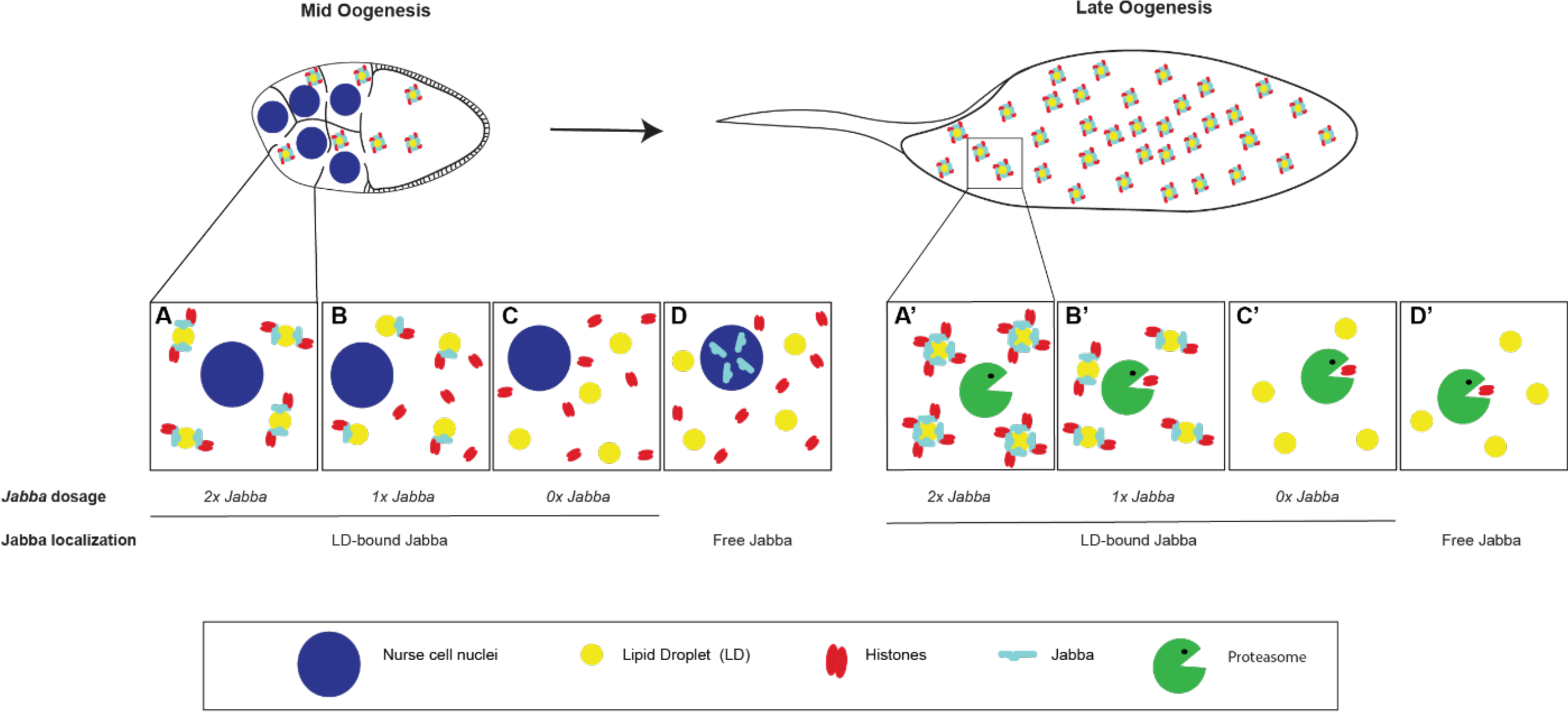
LD association promotes intercellular transport of Jabba to oocytes where it protects histone H2Av from degradation. LD-histone interaction is dependent upon *Jabba* dosage and Jabba localization. In wild-type nurse cells (A), Jabba (cyan) is bound to LDs (yellow) and recruits histone H2Av (red). LDs are transferred to the oocyte. LD-bound Jabba is transferred to the ooplasm and its levels further increase; here Jabba protects H2Av from degradation (A’). Decreasing *Jabba* dosage (*1x Jabba*) reduces the binding capacity of LDs and likely increases the free histone pool in the nurse cell cytoplasm (B). In oocytes, Jabba levels are limiting, leading to partial degradation of the H2Av pool (B’). In *0x Jabba* H2Av is no longer localized to LDs (C, C’). H2Av is unable to evade the degradation machinery and H2Av stores are not maintained in later staged oogenesis (C’). When Jabba is not bound to LDs (Free Jabba), Jabba relocalizes to nurse cell nuclei (D) and is not present in the oocyte and thus cannot prevent H2Av degradation (D’).

Histones H2A and H2B levels in embryos are also Jabba dependent (Li et al., 2012), and the pattern of developmental H2B accumulation during oogenesis (Fig. 2.D) is very similar to that of H2Av (Fig. 2.A-C). We currently do not have a good tool to determine H2A accumulation directly, but predict it will follow the same pattern, since canonical histones are typically regulated in a similar manner (Marzluff et al., 2008). We therefore propose that increasing H2Av, H2B, and H2A accumulation during late oogenesis establishes an LD-bound histone depot for the early embryo.

It will be interesting to determine if the accumulation of histones H3 and H4 follows a similar pattern. On the one hand, as canonical histones, their transcriptional and translational regulation is likely similarly to that of H2A and H2B. However, they are not LD-associated (Cermelli et al., 2006) nor do their embryonic levels depend on Jabba (Li et al., 2012), so there must be mechanistic differences.

### Degradation of excess histones

Newly laid *Jabba ^-/-^* eggs have dramatically reduced levels of histones H2A, H2B, and H2Av compared to the wild type (Li et al., 2012). We find that this divergence in the histone supply is established during late oogenesis, when H2Av and H2B levels rise in the wild type while they drop in the mutant (Fig. 3). Our experiments to partially inhibit proteasome activity reveal that degradation of histones plays a major role in bringing about this difference (Fig. 5, Fig. 9.C-C’). Because high concentrations of the inhibitor arrest development (see also Velentzas et al., 2011), we had to identify intermediate concentrations that allow for egg chambers to develop to stage 14. We infer that under these conditions degradation is only partially inhibited. This partial inhibition presumably explains why H2Av levels in *Jabba* mutants do not rise above the stage 12 level; the residual H2Av turnover, however, is counteracted by ongoing histone synthesis, resulting in the observed stable levels from stage 12 to 14.

In the wild type, proteasome inhibition did not lead to further increase in the H2Av pool beyond levels in untreated egg chambers. This observation was surprising since females with four copies of *Jabba* can accumulate more H2Av than the wild type, which presumably indicates that even the wild type produces some excess H2Av that is degraded if not protected by Jabba. We reason that the direct effects on proteasome turnover are balanced out by indirect effects that compromise histone or Jabba synthesis; indeed, high concentrations of the inhibitor abolish any rise in H2Av levels (Fig.5 - Sup Fig.1).

Turnover of excess histones by the proteasome is well established in yeast (Gunjan & Verreault, 2003). It is one of numerous mechanisms that prevent accumulation of free histones and the resulting cytotoxicity (Bannister & Kouzarides, 2011; Gunjan et al., 2006; Kaygun & Marzluff, 2005; Marzluff et al., 2008). We therefore propose that H2Av turnover in late oogenesis is due to this general protective machinery and that Jabba allows oocytes to continue to deploy this safety feature while also accumulating the extra H2Av needed to provision the early embryo. In this view, Jabba would determine the pool of H2Av to be protected and any excess would be recognized as potential hazard and degraded.

Because the mechanism that targets excess H2Av for proteasomal degradation is not known, we cannot yet test exactly against what type of damage this protective mechanism guards against. In *Drosophila* oogenesis, histone overaccumulation has been achieved by mutating abnormal oocyte (*abo*), a histone-locus specific transcription factor (Berloco et al., 2001). Females homozygous mutant for *abo* lay embryos with mildly (∼1.6x) increased levels of H2B that display a range of phenotypes, from fertilization defects to extra cell cycles to catastrophic early nuclear divisions (Chari et al., 2019; Tomkiel et al., 1995). In yeast, excess of all histones causes wide-spread changes to gene expression, via diverse mechanisms (Singh et al., 2010); in *Drosophila* embryos, excess H2Av results in over-incorporation into chromatin and signs of DNA damage (Johnson et al., 2018; Li et al., 2014). During late *Drosophila* oogenesis, however, the oocyte nucleus is arrested in meiosis with condensed chromatin (Von Stetina & Orr-Weaver, 2011) and minimal transcriptional activity (Navarro-Costa et al., 2016); therefore, it seems unlikely that an oversupply of histones would lead to more incorporation into chromatin or massive transcriptional changes.

Our data suggest that Jabba prevents degradation by physically protecting H2Av. First, H2Av levels scale with *Jabba* dosage, and we can achieve H2Av levels beyond normal by simply doubling the amount of Jabba protein present (Fig. 6.C, D). Second, a version of Jabba that is mislocalized to the nurse cell nuclei and not present in oocytes is unable to support high H2Av levels in oocytes (Fig. 8.D, E, Fig. 9.D, D’). Finally, a *Jabba* mutant unable to bind to H2Av resulted in H2Av levels indistinguishable of those in *Jabba ^-/-^* (Fig. 7.D, E). To further unravel how Jabba prevents degradation, it will be necessary to identify the machinery that targets excess H2Av to the proteasome. Work in *S. cerevisiae* has identified the E3 ligases promoting the turnover of histones H3 and H4 (Singh et al., 2012), but E3 ligases for the other histones remain uncharacterized. An interesting possibility is that Jabba may physically protect H2Av by blocking access to a ubiquitination site. However, Western analysis after MG132 treatment of *Jabba^-/-^* egg chambers did not reveal accumulation of ubiquitinated H2Av bands (data not shown). Though not conclusive, this observation raises the possibility that histone degradation in oocytes occurs independent of ubiquitination. Intriguingly, in mammals, during development of the male germ line, histones are turned over via ubiquitin-independent degradation. (Huang et al., 2016; Khor et al., 2006; Qian et al., 2013).

In embryos, H2Av is not statically bound to Jabba, but constantly exchanges between LDs through the cytoplasm (Johnson et al., 2018). If similar exchange occurs in oocytes, it is not clear how transient interactions with Jabba would be sufficient to prevent proteolytic degradation of H2Av. We speculate that either the transit time through the cytoplasm is negligible relative to the time H2Av spends interacting with Jabba or that H2Av in the cytoplasm is accompanied by a chaperone that also prevents its degradation.

### LD-binding promotes the Jabba availability and, indirectly, of histones

It remains an open question why some histones are stored on LDs, while others (like H3) are apparently stored in the cytoplasm (Cermelli et al., 2006). Our analysis provides a first answer, namely that Jabba, the protein necessary to stabilize the oocyte pool of H2A, H2B, and H2Av, gets trapped in nurse cell nuclei if it is unable to bind to LDs. As a result, it is absent from the oocyte and thus cannot perform its protective function there. We propose that LD binding ensures proper intercellular transport of Jabba, thereby safeguarding its ability to function in the ooplasm (Fig. 9). In the embryo, the association of H2Av with LDs has a distinct function. Here, transient sequestration of H2Av retards its import into the nucleus and thus prevents H2Av overaccumulation inside the nucleus and on chromatin (Johnson et al., 2018).

The ooplasmic stores of histones H3 and H4 are presumably also bound to some partner protein that keeps them soluble and protects them from degradation, perhaps the histone chaperone NASP (Campos et al., 2010; Finn et al., 2012) . As these histones are apparently cytoplasmic, it is not clear how they and their binding partners avoid being trapped in nurse cell nuclei. Maybe synthesis of H3 and its binding partner only ramp up in the oocyte, after dumping. We find some evidence that such delayed translation may also be true for Jabba; the ooplasmic synthesis of Jabba accounts for about half of the total Jabba present at stage 14 (Fig.7 - Sup. Fig. 1A, B). As final histone storage is dependent on *Jabba* dosage, this synthesis of Jabba in oocytes may have to be supplemented by a pool that is made earlier in the nurse cells in order to achieve the required levels. Alternatively, H3 and its binding partners may display transient interactions with some cytoplasmic organelle that supports their transfer from nurse cells to oocytes, but this interaction is lost by early embryonic stages, as there H3 behaves like a cytoplasmic protein (Cermelli et al., 2006; Shindo & Amodeo, 2019).

It is also not clear why histones are stored on LDs rather than some other cytoplasmic structure. A priori, any cytoplasmic organelle that is transferred to oocytes could suppress the import of the histone anchor into nurse cell nuclei and promote transport into the oocyte. It is possible that the large surface area of the hundreds of thousands of LDs provides a readily available and, at these developmental stages, metabolically inert platform for recruitment. Other organelles may not have enough storage capacity or might be functionally impaired by massive amounts of positively charged proteins on their surface. Alternatively, LD association may be an evolutionary accident and many other organelles might in principle be able to store histones. Our identification of a region of Jabba that fails to localize to LDs but is sufficient for histone binding (Jabba ^HBR^; Fig. 8) will make it possible to address this question in the future, by transplanting Jabba ^HBR^ to other organelles and test the consequences for histone storage and normal developmental progression.

But why is Jabba ^HBR^ mislocalized to nurse cell nuclei at all? We suspect that it is dragged along when histones are imported into those nuclei and/or it is retained in nuclei because it binds to the histones present in nurse cell chromatin. In either model, it would be Jabba ^HBR^’s histone binding ability that leads to its mislocalization. As the ability to bind histones is important for Jabba ^HBR^ to protect them from degradation, mislocalization cannot be avoided unless Jabba ^HBR^ is somehow anchored in the cytoplasm. We have not yet been able to test this idea directly as a Jabba ^HBR^ unable to interact with histones still accumulates in the nucleus, in cultured cells as well as in nurse cells (unpublished observations). We hypothesize that a cryptic nuclear localization signal in Jabba ^HBR^ promotes nuclear transport even in the absence of histone binding. This second pathway currently masks what contribution histone binding makes to nuclear accumulation/retention of Jabba ^HBR^.

### Broader insights into protein sequestration

Our analysis of histone sequestration in *Drosophila* oogenesis may also shed light on how LDs regulate other proteins. It is now well-established that LDs can transiently accumulate proteins from other cellular compartments (Welte, 2007). This has been particularly documented for a number of proteins involved in nuclei acid binding/transcriptional regulation: For example, in mammalian cells, MLX and Perilipin 5 can either be present on LDs or move into the nucleus to regulate transcription (Gallardo-Montejano et al., 2016; Mejhert et al., 2020). In bacteria, the transcriptional regulator MLDSR (microorganism lipid droplet small regulator) is sequestered to LDs under stress conditions (Zhang et al., 2017). Finally, the core protein of Hepatitis C virus (HCV) is transiently recruited to LDs before is it used in viral assembly (Filipe & McLauchlan, 2015). Additional examples of proteins with roles in /from other compartments have also been described (Welte & Gould, 2017). Like for Jabba and H2Av, LD association of such “refugee proteins” (Welte, 2007) may prevent their premature turnover or may promote their delivery to the correct intra-or intercellular location. Indeed, in the absence of LD binding, HCV core is degraded (Beller et al., 2006; Camus et al., 2013; McLauchlan et al., 2002). In addition, a role for LDs in protein delivery to distant cellular compartments could be particularly important to studies in neurons. Recent discoveries indeed suggest important, but largely uncharacterized roles for LDs in this cell type (Pennetta & Welte, 2018; Wat et al., 2020).

## Materials and Methods

### Fly Stocks

Oregon R was used as the wild-type strain. Alleles *Jabba ^DL^* and *Jabba ^zl01^* were used to analyze ovaries and embryos lacking *Jabba.* (Li et al., 2012). To analyze embryos/ovaries with varying *Jabba* levels, these additional genotypes were used: Df(2R)Exel7158/*CyO* (Bloomington *Drosophila* Stock Center (BDSC: 7895; FLYB: FBab0038053)) carries a large deletion that encompasses *Jabba* and is used to reduce Jabba dosage; for simplicity it is referred to as Jabba ^DF^ in the figures. *Jabba ^low^* was derived from *Jabba ^d03001^* (a PBac insertion, P[XP]d03001, obtained from The Exelixis Collection at Harvard Medical School) by imprecise P-element excision. To generate *4x Jabba*, *gJabba* (previously described in Johnson et al., 2018) was introduced into an otherwise wild-type background. H2Av dosage was increased using the genomic transgene *gH2Av* (Johnson et al., 2018) which expresses H2Av under endogenous control at normal levels. The following fluorescently-tagged histone stocks were used to obtain H2Av-RFP, H2Av-GFP and H2B-mEos3.2 ovaries with varying *Jabba* dosage: H2Av-RFP Bloomington *Drosophila* Stock Center (BDSC: 23650; FLYB: FBst0023650), H2Av-GFP Bloomington *Drosophila* Stock Center (BDSC: 24163; FLYB: FBst0024163), H2B-mEos3.2, a gift from Michael Eisen (Mir et al., 2018). The *LSD-2* null allele was previously described (Welte et al., 2005). *Jabba B FLAG-HA* expresses Jabba B with a C-terminal FLAG tag under UAST control and was generated by the *Drosophila* Protein interaction Map consortium (Guruharsha et al., 2012) and obtained from the Bangalore Fly Facility. The generation of the following mCherry tagged lines is described below: *Jabba B ^FL^*, *Jabba B ^del aa 228-243^*, and *Jabba ^HBR^*. P{mat4-GAL-VP16} V2H was used to drive expression using the Gal4/UAS system (Bloomington *Drosophila* Stock Center (BDSC: 7062; FLYB: FBst0007062)).

### Transgene generation/Molecular Biology

The Jabba ^HBR^ construct (amino acids 1-192) was generated using the oligonucleotides JabbaB192_321forward: 5’-CAGGGTTTAAGCAATTTCGTAGT and Vector_Start_re: 5’-P-CATGGTGGCGGCCGCGGAGC using standard in vitro mutagenesis protocols and the JabbaB full length construct as template. GFP or luciferase tagged versions of the proteins were generated using the Gateway recombination cloning procedure (Invitrogen, Carlsbad, CA) and custom-made vectors.

To generate mCherry tagged Jabba B ^FL^, Jabba B del ^aa 228-243^, and Jabba ^HBR.^ transgenes, the desired sequences were amplified from cDNA samples and donor plasmids (Kolkhof et al., 2017), and then cloned into the pENTR Gateway plasmid system (Invitrogen). The Entry plasmids were then recombined using the Gateway recombination cloning system (Invitrogen) into pPTAPmChW attB (described in Hudson & Cooley, 2010) destination plasmids. Transgenic lines were created by BestGene Inc. (Chino Hills, CA) using PhiC31 integrase-mediated transgenesis. All insertions were incorporated onto the third chromosome at site 68A4.

### Western Analysis

Egg chambers/ embryos were heat-fixed in 1x Triton Salt Solution, (embryos were devitellinized), sorted by stage, and boiled in Laemmli buffer (Bio-RAD, Hercules, CA) for 15 minutes. Proteins were separated using 4-15% SDS-PAGE gels (Bio-RAD) and transferred to PVDF membranes (Immobilon-FL, EMD Millipore, Burlington, MA). Transfers were performed in N-cyclohexyl-3-aminopropanesulfonic acid (CAPS) buffer (10mM CAPS, 10% Methanol) at 50V for 40 mins (Jabba) or Towbin buffer (10% 10x Tris-Glycine solution, 20% Methanol) at 80V for 30 mins (HA and H2Av). Immunodetection was done using the following primary antibodies – rabbit anti-Jabba (1:5000) (previously described in Johnson et al., 2018), rabbit anti-H2Av (1:2500) (Active Motif, Carlsbad, CA), rat anti-HA (1:1000) (Clone 3F10; Roche Diagnostics, Indianapolis, IN) and mouse anti-α-Tubulin (1:10,000) (Cell Signaling, Danvers, MA) and secondary antibodies – IRDye® 800CW Goat anti-Rabbit IgG, IRDye® 680RD Goat anti-Mouse IgG, and IRDye® 680RD Goat anti-Rat IgG (1:10,000) (Li-COR, Lincoln, NE). The same membranes were probed with the indicated primary antibody and mouse anti-α-tubulin as a loading control. Images were acquired using Li-COR Odyssey CLx and fluorescence was quantified using Image StudioLite Software.

### Immunostaining and Staining

Embryos were collected on apple juice agar plates and then de-chorionated with 50% bleach. *In vivo* centrifugation was then performed (Tran & Welte, 2010) and embryos were subsequently fixed with 4% formaldehyde/1x Phosphate Buffered Saline (PBS) for 20 minutes. Embryos were devitellenized using heptane/methanol and subsequently washed in 1x PBS/0.1%Triton X-100.

Ovaries were dissected from females maintained on dry yeast at 25 °C overnight. Samples were then fixed with 4% formaldehyde/1x Phosphate Buffered Saline (PBS) for 15 minutes and subsequently washed in 1xPBS/0.1%Triton X-100.

Embryo and ovary samples were then sorted, counted (to ensure equal numbers of samples were prepped) and blocked over night at 4°C in 10% BSA/0.1Triton/1x PBS solution. Incubation of primary antibodies (Jabba, H2Av) were performed at a 1:1000 concentration overnight. Samples were washed and probed with secondary antibodies at a final concentration of 1:1000 (goat anti-rabbit Alexa 488, goat anti-rabbit Alexa 594, goat anti-mouse Alexa 488). For DNA staining, embryos/egg chambers were stained using 4′,6-diamidino-2-phenylindole (DAPI) (Sigma-Aldrich, St Louis, MO). Lipid droplets were stained using either Nile Red (Sigma-Aldrich) or BODIPY™ 558/568 C12 (4,4-Difluoro-5-(2-Thienyl)-4-Bora-3a,4a-Diaza-s-Indacene-3-Dodecanoic Acid) (Thermo Fisher Scientific, Waltham, MA).

### Microscopy/Imaging

*In vivo* centrifugation was performed as described by Tran *et al.,* 2010 (Tran & Welte, 2010). Centrifuged embryos were imaged with a Nikon Eclipse E600 epifluorescence microscope, using an 20X objective. For live imaging, ovaries were dissected, from flies maintained on dry yeast overnight at 25 °C. Egg chambers were placed on a coverslip and overlaid with Oil 10 S, VOLTALEF (VWR Chemicals, Radnor, PA). Imaging of both live and fixed samples was performed using a Leica SP5 confocal microscope. All images were assembled using Abode Illustrator.

### RNA Extraction and qPCR

Stage 14 stage egg chambers were dissected from females maintained on dry yeast at 25 °C for 48 hours. Unfertilized eggs were collected on apple juice plates and de-chorionated in 50% bleach. Trizol ^TM^ Reagent (Invitrogen) was used according to the manufacturer’s instructions to extract total RNA. Recovered RNA was measured to ensure that equal concentrations of RNA samples were digested with DNAse (Thermo Fisher Scientific). cDNA was then synthesized using Maxima H Minus First Strand cDNA Synthesis kit (Thermo Fisher Scientific). To estimate the H2B/H2A/H2Av mRNA levels, Power SYBR Green PCR Master Mix (Thermo Fisher Scientific) was used and qPCR was performed on a StepOnePlus Real time PCR System (Applied Biosystems, Foster City, CA). Primers for qPCR:

H2A Fw:5’-CGCAACGACGCGGCGTTAAA-3’,Rv:5’-GCCTTCTTCTCGGTCTTCTTG-3’

H2Av Fw: 5’-GTGTACTCCGCTGCCATATT-3’, Rv: 5’-CGAATGGCGAGCTGTAAGT-3’,

H2B Fw: 5’-CCATCCTGACACCGGAATTT-3’, Rv: 5’-TCGAGCGCTTGTTGTAGTG-3’

H2B/H2A/H2Av mRNA levels were normalized to the reference gene 60S Ribosomal ProteinL24 (RPL24).

Fw: 5’-TTTCTACGCCAGCAAGATGAAAA-3’, Rv: 5’-ATTGCGCTTCATCAGGTAGG-3’

### Triglyceride (TAG) Assays

Triglyceride amounts were measured using approaches adapted from Palanker *et al*. (Palanker et al., 2009). 100 egg chambers were homogenized in 100 μl PBS, 0.5% Tween 20 and immediately incubated at 70°C for 5 min. 20 μl of the homogenate was incubated with either 20 μl PBS or 20 μl Triglyceride Reagent (Sigma-Aldrich) for 30 min at 37°C. The samples were then centrifuged using an Eppendorf Microcentrifuge (Model 5415D) at maximum speed for 3 minutes. Then 30 μl of each sample was transferred to a 96-well plate and incubated with 100 μl of Free Glycerol Reagent (Sigma-Aldrich) for 5 min at 37°C. Samples were assayed using a SpectraMax M2e spectrophotometer at 540 nm. TAG levels were determined by subtracting the amount of free glycerol in the PBS-treated sample from the total glycerol present in the sample treated with Triglyceride reagent. TAG amounts were normalized to the total protein amounts in each sample using a Pierce ^TM^ BCA Protein Assay Kit (Thermo Fisher Scientific). The data were analyzed using a Student’s t test.

### Polysome Analysis

Ovaries from flies (maintained on dry yeast paste overnight at 25°C) were dissected in PBS. Stage 14 egg chambers were treated with 10 μg/ml puromycin dihydrochloride from *Streptomyces alboniger* (Millipore Sigma, St. Louis, MO) in PBS on ice. Egg chambers were then lysed [lysis buffer: 20 mM Tris-HCl, 140 mM KCl, 5 mM MgCl2, 0.5 mM Dithiothreitol (DTT), 1% Triton, 0.1 mg/ml cycloheximide (Sigma-Aldrich), 1mg/ml Heparin, 50 units/ml RNAsin (Applied Biosystems)] and equal amounts of lysate were cleared by centrifugation at 10,000 rpm for 5 min at 4°C. Sucrose sedimentation profiles were performed as described (Connolly et al., 2008; Maki et al., 2002), onto a 10-50% sucrose gradient [containing 50 mM Tris-HCl (pH 7.8), 10 mM MgCl2, 100 mM KCl, 2 mM DTT, 100μg/ml cycloheximide, 10-50% sucrose]. Samples were spun using a Beckman Coulter Optima ^TM^ L-90K Ultracentrifuge in an SW41 rotor (36,000 rpm) for 2.5 hours at 4° C. Gradients were analyzed using a Biocomp Piston Gradient Fractionator with a BIORAD Econo UV Monitor with a Full Scale of 1.0. Data was recorded using DataQ DI-158-UP data acquisition software. Samples were collected from fractions containing the 80S peak and polysome peaks. RNA was then extracted using Trizol ^TM^ Reagent.

### In *vitro* egg maturation (IVEM)

IVEM experiments were adapted from Spracklen *et al.,* 2013 (Spracklen & Tootle, 2013). *Drosophila* females were fed dry yeast overnight and maintained at 25 °C. Ovaries were dissected into freshly prepared IVEM media (1x Schneider’s *Drosophila* media (Gibco, Thermo Fisher Scientific), heat-inactivated 10% fetal bovine serum (Invitrogen) and 1x penicillin/streptomycin glutamine (100x, Gibco, Thermo Fisher Scientific)). Individual ovarioles were isolated so that morphological stages of egg chamber development were easily distinguishable. Egg chambers of interest were then transferred into a mutli-well plate with 500 μl fresh media with the addition of MG132 (EMD Millipore) or DMSO. Plates were placed in the dark for *in vitro* development. After 6 hours, egg chambers were mounted in Oil 10 S, VOLTALEF (VWR Chemicals) for live imaging.

### General tissue culture procedures

*Drosophila* Kc167 cells were cultured in Schneider’s *Drosophila* medium with L-glutamine (PAN Biotech, Aidenbach, Germany) including 10% heat-inactivated fetal calf serum (PAN Biotech) and Penicillin/Streptomycin according to general culturing procedures.

For microscopy, cells were transferred to cover slips, fixed with 4% paraformaldehyde, counterstained with Hoechst33342 for DNA, and embedded with Mowiol. Images were recorded using a Zeiss LSM710 confocal microscope.

### Luciferase complementation assay

Protein-protein interactions were detected using a split-luciferase complementation assay as described previously (Kolkhof et al., 2017). In brief, putative interactors were tagged at the N- or the C-terminus with either the N-or C-terminal half of the Gaussia princeps luciferase enzyme (termed “hGluc(1)” or “hGluc(2)” fragments). Cells were seeded in transparent tissue-culture 96-well plates (Sarstedt, Thermo Fisher Scientific) and allowed to adhere overnight. The next day, plasmids encoding putative interactors were transfected in triplicate using the Effectene transfection reagent (Qiagen, Hilden, Germany) and a final concentration of 400µM BSA-bound oleic acid were added. After 4 days, the medium was removed from the cells, and the cells were – after one PBS washing step – frozen at -80°C to facilitate homogenization. Lysis and luciferase activity detection was performed using the Gaussia-Juice Luciferase Assay (PJK Biotech, Kleinblittersdorf, Germany) according to the manufacturer’s description in white opaque 96-well plates (OptiPlate, PerkinElmer, Waltham, MA) and using a Synergy Mx microplate reader (BioTek, Winooski, VT). A part of the lysate was kept and utilized to measure the protein content using a standard BCA protein assay (Pierce).

As controls, each assay included untransfected cells (baseline based on the background luciferase activity) as well as cells transfected with plasmids encoding the yeast GCN4 leucine zipper protein fused to hGluc(1) as well as hGluc(2), which is known to dimerize.

### Quantitation and Image Analysis

Image J was used to quantitate all images.

To determine the Mean Fluorescence Intensity (AU) of histones, the fluorescence intensity of RFP, GFP, or mEos3.2 (in a constant ROI) was measured. Because fluorescent intensity in nurse cells was quite variable from egg chamber to egg chamber, we chose not to quantify the signal in nurse cells and concentrated on oocytes. Oocytes were imaged just below the follicle nuclei, where LDs were most abundant. To determine the area covered by lipid droplets, confocal images of egg chambers, imaged using confocal reflection microscopy (CRM) (excitation 633 nm/emission 623-643 nm) (Gaspar & Szabad, 2009) were analyzed. The boundaries of the area covered by lipid droplets was identified via thresholding and the area was measured. This analysis was repeated for egg chambers stained with BODIPY™.

For the IVEM experiments in Fig. 5, the average mean fluorescence intensity (AU) of *WT; H2Av-RFP* was calculated. To obtain each data point for the ratio of *Jabba ^-/-^*: WT, individual *Jabba ^-/-^*; *H2Av-RFP* mean fluorescence intensity (AU) was divided by the previously calculated average mean fluorescence intensity (AU) of *WT; H2Av-RFP*. This was repeated for each treatment condition.

Statistics were performed using unpaired Student’s t test or ANOVA followed by a Tukey test. (Prism7, GraphPad) All graphs were generated using Prism 7.

To determine the expression levels of Jabba B and Jabba B ^del aa 228-243^, embryos were immunostained using anti-Jabba antibody and then imaged by confocal microscopy. For quantitation of anti-Jabba signal (Fig. 7 - Sup Fig. 1.G), the embryo boundary was determined using thresholding and image calculator tool in Image J. The anti-Jabba signal in the embryo was measured. The fluorescence intensity of the embryo was then measured by subtracting the background signal in a region outside the embryo (using the original image) from the anti-Jabba signal of the embryo. Summed fluorescence intensity was then divided by the area of the embryo (Fig. 7 - Sup Fig. 1.H). To quantify Jabba levels of Jabba B and Jabba B ^del aa 228-243^, the fluorescence intensity/area of Jabba B and Jabba B ^del aa 228-243^ was compared to the fluorescence intensity/area of embryos expressing varying levels of endogenous Jabba (The expression levels of these embryos were determined by Western analysis in Fig. 7 - Sup Fig. 1.E,F). Using this comparison, embryos of similar Jabba expression patterns were identified. This method of quantitation was employed to avoid the potential for differential transfer endogenous Jabba, and mCherry-tagged constructs, given their vast disparity in apparent molecular weight.

## Acknowledgements

We thank Michael Eisen, the Bloomington Stock Center, the Exelixis *Drosophila* Collection at Harvard Medical School and the Bangalore Fly Facility for fly stocks. We are grateful to Sina Ghaemmaghami for discussion and advice and to Xin Bi, Daniel Bergstralh, and Jeffrey Hayes for comments on the manuscript. We also thank Tina Tootle and Roxanne Keltz for technical assistance.

