## Supplemental figures for "Sequestration to lipid droplets promotes histone availability by preventing turnover of excess histones"

#### Figure 5 Supplemental Figure 1

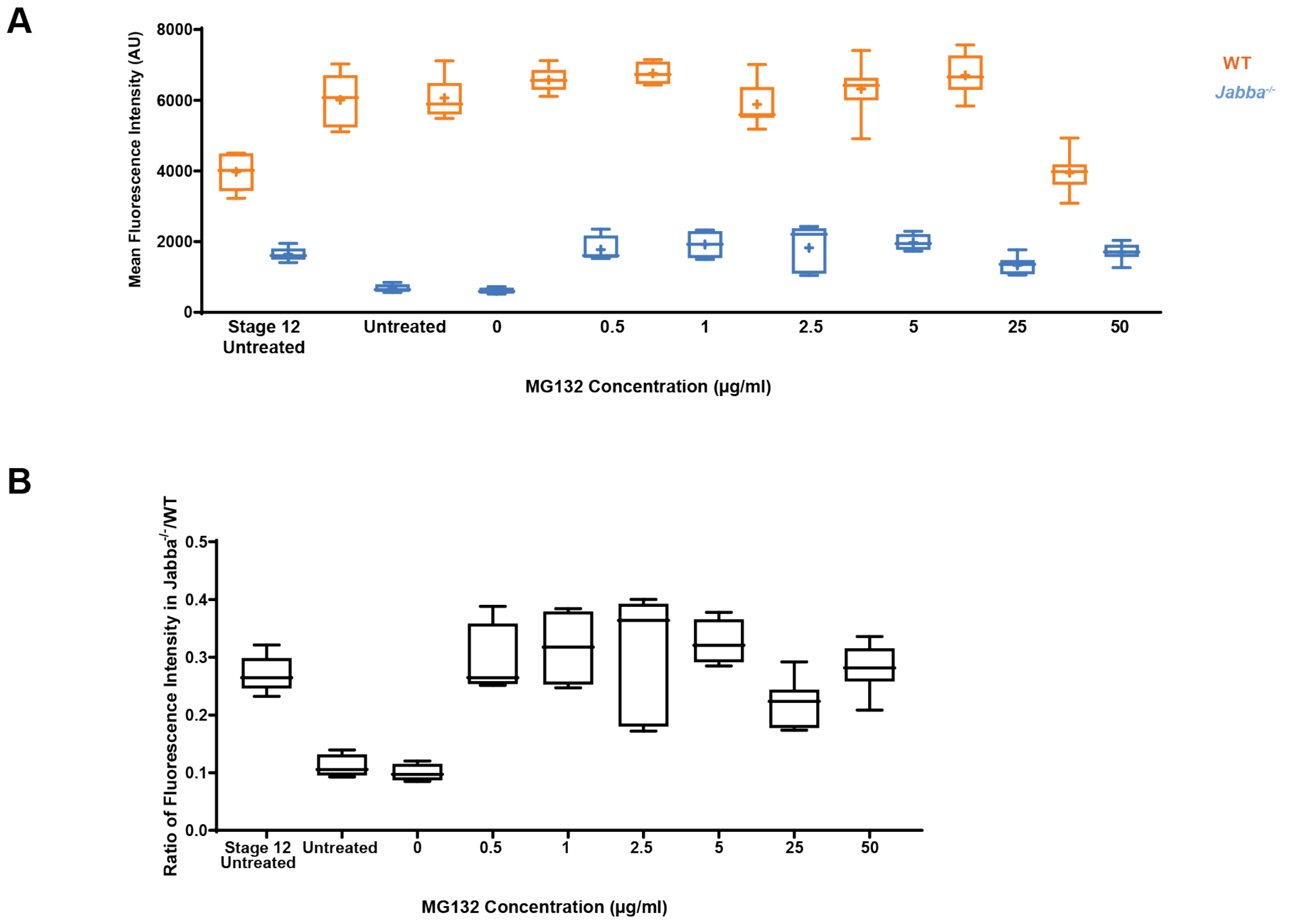

**Figure 5-Supplemental Figure 1: H2Av degradation in the absence of Jabba**

A) Mean fluorescence intensity (AU) in stage 14 WT (orange) or *Jabba* $^{-/-}$ (blue) egg chambers after IVEM. Error bars represent SD. Length of box plot represents the 25$^{th}$ and 75$^{th}$ percentile. Line represents median, cross indicates the mean. (B) The ratio of the average mean fluorescence intensity (AU) of *Jabba* $^{-/-}$*:* WT for indicated concentrations of MG132 treatment. Stage 12 Untreated = stage 12 egg chambers after dissection; Untreated = *in vitro* culture in IVEM media; 0, 0.5, 1, 2.5, 5, 25, 50 µg/ml= indicates the various concentrations of MG132 (diluted in DMSO) used in IVEM media. A subset of these data is presented in Fig. 5B.

### Figure 7 Supplemental Figure 1

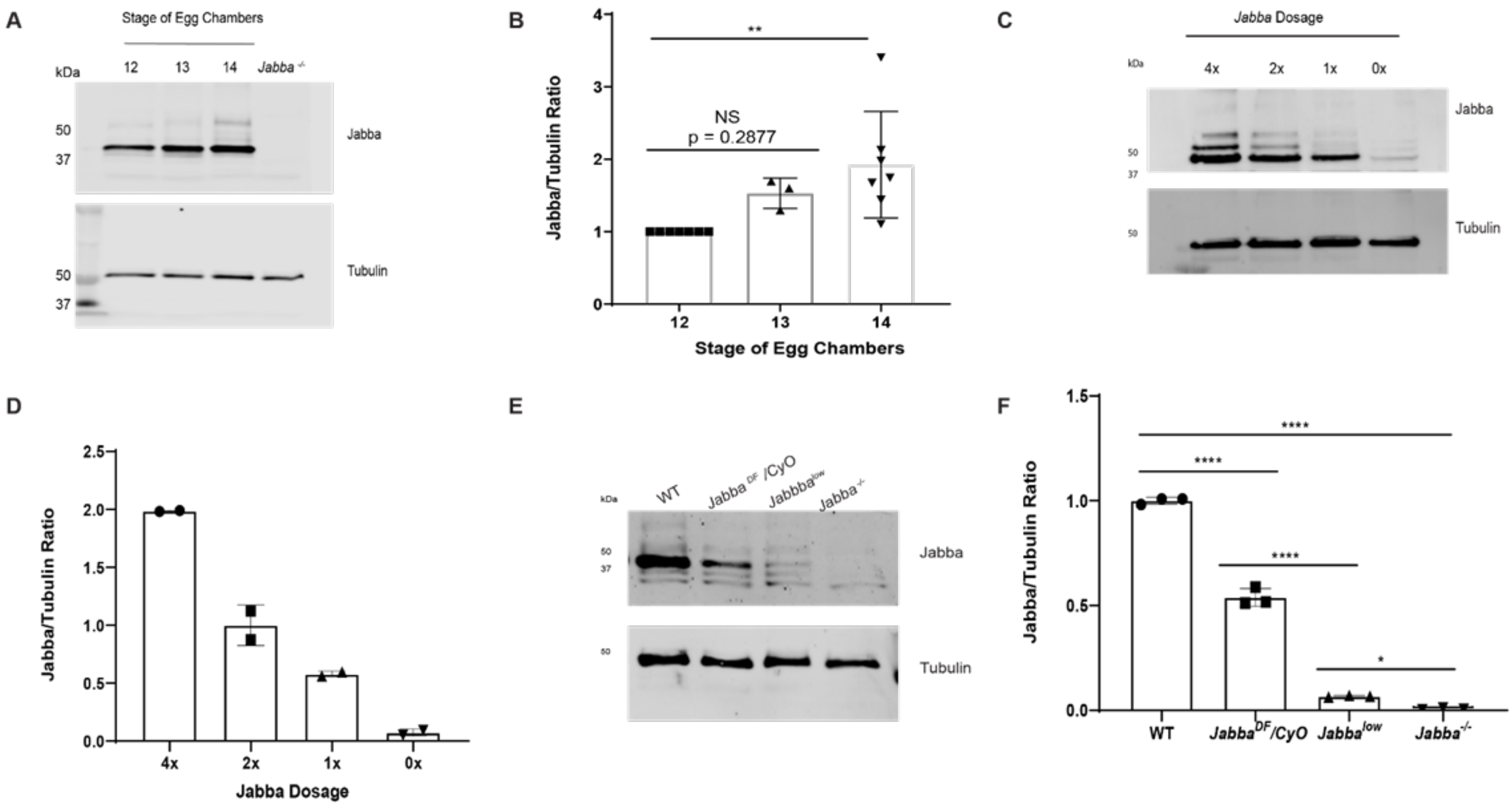

G

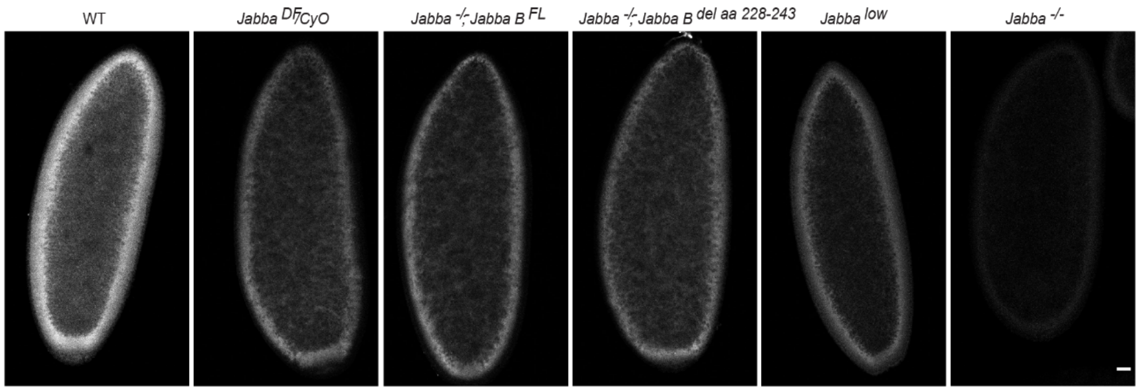

WT
Jabba DF/CyO
Jabba -/-; Jabba B FL
Jabba -/-; Jabba B del aa 228-243
Jabba low
Jabba -/-


H

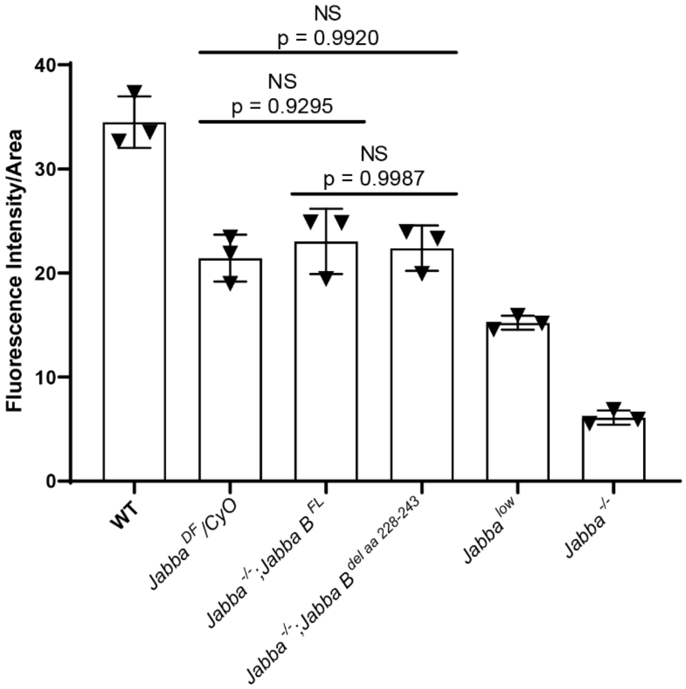

NS
p = 0.9920
NS
p = 0.9295
NS
p = 0.9987
Fluorescence Intensity/Area
40
30
20
10
0
WT
Jabba DF /CyO
Jabba -/- ;Jabba B FL
Jabba -/- ;Jabba B del aa 228-243
Jabba low
Jabba -/-

**Figure 7-Supplemental Figure 1: Analysis of Jabba levels during oogenesis and embryogenesis.**

A) Western analysis of Jabba levels in egg chambers from stage 12 to 14. *Jabba* $^{-/-}$ was used as a negative control. Membranes were probed for Jabba and for tubulin, as a loading control. (B) Quantitation of (A) expressed as the Jabba/tubulin ratio normalized to stage 12. N = 3-7. p values were calculated using one-way ANOVA followed by Tukey's test. Stage 12 vs Stage 13, p = 0.2877; Stage 12 vs stage 14, p = 0.0085. (C) Jabba expression scales with *Jabba* dosage. Western analysis of Jabba levels in stage 14 egg chambers of flies expressing varying dosages of *Jabba*. Membranes were probed for Jabba and for tubulin, as a loading control. (D) Quantitation of (C) expressed as the Jabba/tubulin ratio normalized to 2x Jabba. N = 3. E) Anti-Jabba Western of embryos from wild-type flies and flies of various *Jabba* genotypes. *Jabba* $^{low}$ expresses low levels of wild-type Jabba. (F) Quantitation of (E) expressed as the Jabba/tubulin ratio normalized to wild type. In *Jabba* $^{DF}$*/CyO*, Jabba is detected at roughly half the expression of wild type. Jabba protein is decreased in *Jabba* $^{low}$. N = 3. p values were calculated using one-way ANOVA followed by Tukey's test. WT vs Jabba $^{-/-}$, p <0.0001; WT vs *Jabba* $^{DF}$*/CyO*, p <0.0001; *Jabba* $^{DF}$*/CyO* vs *Jabba* $^{low}$, p <0.0001; *Jabba* $^{low}$ vs *Jabba* $^{-/-}$, p = 0.0485. (G) Anti-Jabba (white) immunostaining of embryos of various *Jabba* genotypes. Scale bar represents 25 µm. (H) Quantitation of Fluorescence Intensity/Area for embryos in (G). The Fluorescence Intensity/Area for *Jabba* $^{-/-}$*; Jabba B* $^{FL}$ and *Jabba* $^{-/-}$*; Jabba B* $^{del\ aa\ 228\text{-}243}$ embryos is comparable to *Jabba* $^{DF}$*/CyO* embryos (expressing 1x copy of *Jabba*). p values were calculated using a one-way ANOVA followed by Tukey's test. WT vs *Jabba* $^{-/-}$*; Jabba B* $^{FL}$, p = 0.0003; *Jabba* $^{DF}$*/CyO* vs

*Jabba* $^{-/-}$*; Jabba B* $^{FL}$, $p = 0.9295$; *Jabba* $^{DF}$*/CyO* vs *Jabba* $^{-/-}$*; Jabba B* $^{del\ aa228\text{-}243}$; $p = 0.9920$; *Jabba* $^{-/-}$*; Jabba B* $^{FL}$ vs *Jabba* $^{-/-}$*; Jabba B* $^{del\ aa228\text{-}243}$; $p = 0.9987$; WT vs *Jabba* $^{DF}$*/CyO*, $p < 0.0001$

**Figure 7 Supplemental Figure 2**

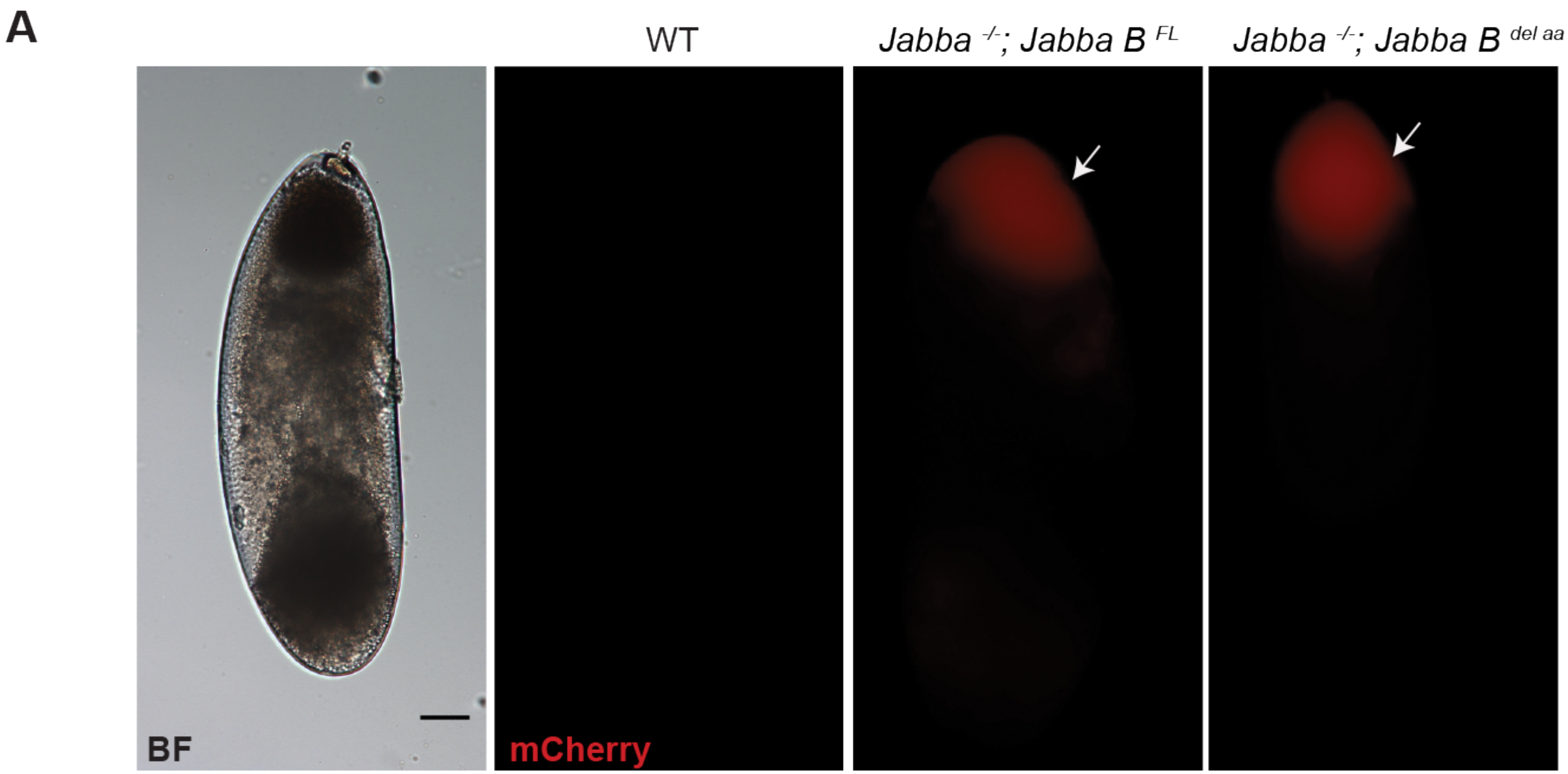

**Figure 7-Supplemental Figure 2: Jabba B $^{FL}$ and Jabba B $^{del\ aa\ 228\text{-}243}$ can associate with LDs.**

A) mCherry signal (red) is detected in the LD layer (arrow) of centrifuged embryos expressing the Jabba transgenes, but not in wild-type embryos. BF = Brightfield image of centrifuged wild-type embryo. Scale bar represents 50 μm.

**Figure 8 Supplemental Figure 1**

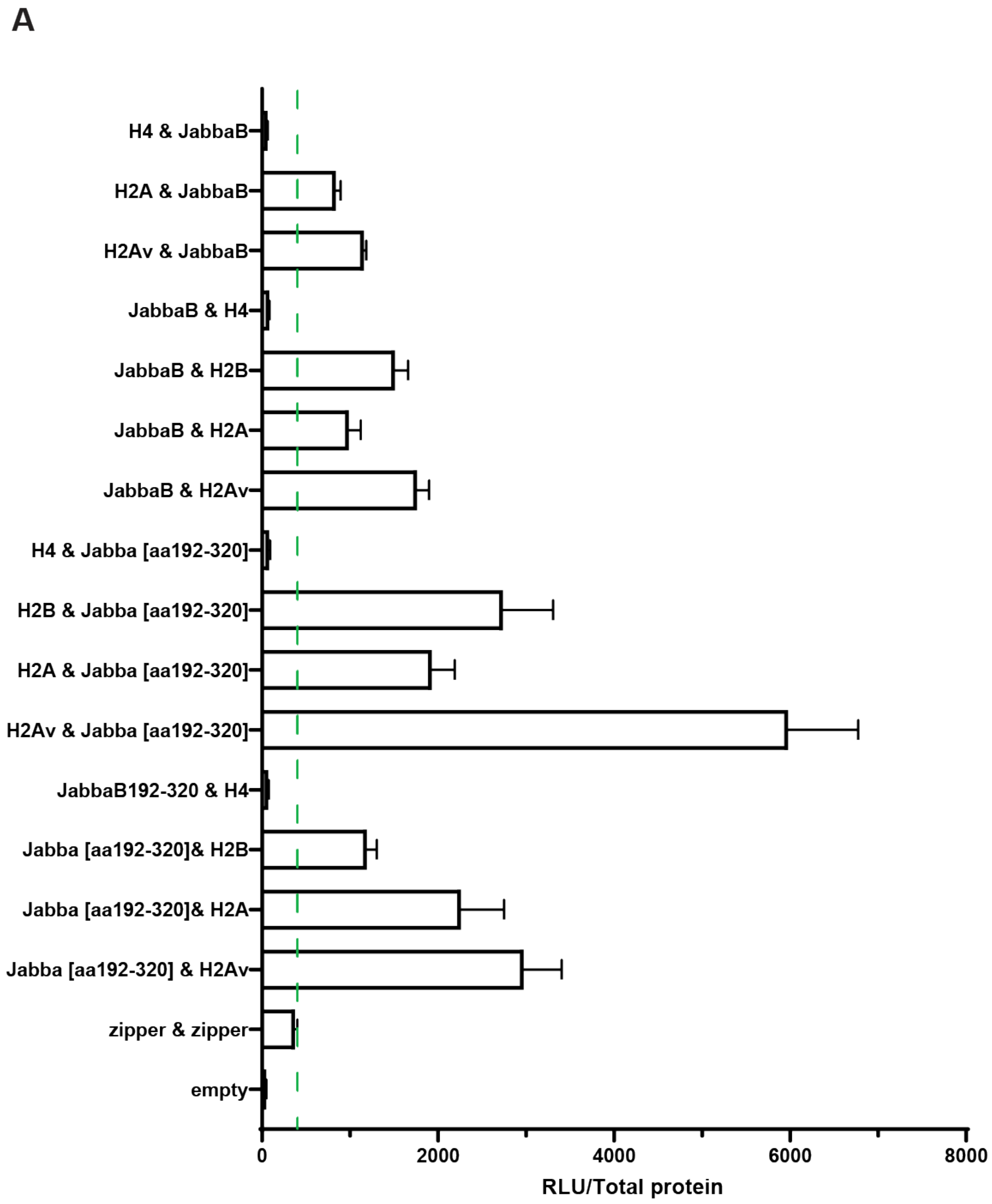

**Figure 8-Supplemental Figure 1: Split luciferase complementation assay showing results for the co-expression of the indicated proteins.**

Luciferase complementation readings are expressed as relative light units (RLU) per µg total protein (RLU/Total protein). Error bars represent standard error of the mean (SEM). The green dashed line represents the “zipper-zipper” control, which was used as a threshold for positive interactions. The fusion proteins were tagged with either the N-or C-terminal half of the *Gaussia princeps* luciferase enzyme (termed “hGluc(1)” or “hGluc(2)” fragments, respectively). Proteins are denoted in the graph as hGluc(1) and hGluc(2). A subset of these data is shown in Fig. 8B.
